# Visuomotor prediction during action planning in the human frontoparietal cortex and cerebellum

**DOI:** 10.1101/2023.07.20.549926

**Authors:** Felix Quirmbach, Jakub Limanowski

## Abstract

The concept of forward models in the brain, classically applied to describing on-line motor control, can in principle be extended to action planning; i.e., assuming forward sensory predictions are issued during the mere preparation of movements. To test this idea, we combined a delayed movement task with a virtual reality based manipulation of visuomotor congruence during functional magnetic resonance imaging (fMRI). Participants executed simple hand movements after a delay. During the delay, two aspects of the upcoming movement could be cued: the movement type and the visuomotor mapping (i.e., (in)congruence of executed hand movements and visual movement feedback by a glove-controlled virtual hand). Frontoparietal areas showed increased delay period activity when preparing pre-specified movements (cued > uncued). The cerebellum showed increased activity during the preparation for incongruent > congruent visuomotor mappings. The left anterior intraparietal sulcus (aIPS) showed an interaction effect, responding most strongly when a pre-specified (cued) movement was prepared under expected visuomotor incongruence. These results suggest that motor planning entails a forward prediction of visual body movement feedback, which can be adjusted in anticipation of nonstandard visuomotor mappings, and which is likely computed by the cerebellum and integrated with state estimates for (planned) control in the aIPS.

## Introduction

Our current understanding of adaptive motor control largely builds upon the concept of internal forward models that predict the sensory consequences of one’s movement based on motor signals (Miall & Wolpert, 1996; Wolpert & Kawato, 1998; Shadmehr, & Krakauer, 2008). This forward prediction can be used to calculate a dynamic estimate of the effector’s state, enabling ‘on-line’ control despite sensory delays (Todorov & Jordan, 2002; Wolpert & Ghahramani 2000). Moreover, forward predictions can be updated by sensory prediction errors, enabling, for instance, the adaptation of control to new sensorimotor mappings (e.g., altered visual movement feedback, Tseng et al., 2007; Grafton et al., 2008; Shadmehr et al., 2010). The respective computations have been linked to the posterior parietal cortex (PPC) and the cerebellum (Miall et al. 1993; Wolpert et al. 1998; Blakemore et al., 1998; Kilteni et al., 2023), and their interaction with (pre)motor areas and the basal ganglia implementing the task and control policies (Shadmehr & Krakauer, 2008; Scott, 2012; Parr et al., 2021). Thus, activity of the PPC and cerebellum has been associated with the detection of ‘nonstandard’ visuomotor mappings (Leube et al., 2003; Nahab et al., 2011; Limanowski et al., 2017; Arikan et al., 2019), and with the learning process required to adapt motor control to those mappings (Ogawa et al., 2006; Izawa et al., 2012; Van Kemenade et al., 2017, 2019; Ohata et al., 2020; Kufer et al., 2024). In line with this, brain stimulation and clinical case studies have linked the efficacy of sensory forward predictions directly to intact PPC and cerebellar function (Desmurget et al., 1999; Synofzik et al., 2008; Parr et al., 2021; Schmitter & Straube, 2024).

Besides on-line control and learning, however, the internal forward model framework can, in principle, also be applied to action *planning* (Stanley & Miall, 2009; Scott, 2012); in line with the proposed functional equivalence of planning and overt action control in the brain (Jeannerod & Decety, 1995; Lotze, 1999). A derivative hypothesis of this extension is that the motor system should also issue sensory predictions when merely planning actions; i.e., in the absence of overt action and associated sensory movement feedback (Kuang et al., 2015; Pilacinski et al., 2018; Kilteni et al., 2018).

We tested this idea by combining a delayed movement paradigm with a virtual reality based manipulation of visuomotor mapping inside an MR scanner. In delayed movement tasks, movements are pre-cued (Rosenbaum, 1980); neural activity during the delay period is thought to encode specific aspects of motor planning, such as the selection of effectors or goals, and the preparation of the appropriate control policy and motor commands (Crammond & Kalaska, 1989; Hoshi & Tanji, 2002; Fernandez-Ruiz et al., 2007; Bernier et al., 2012; Ariani et al., 2022). These planning processes have been linked to activity in the dorsal premotor cortex (PMd) and the PPC (Churchland et al., 2010; Cisek & Kalaska, 2005; Ferraina & Bianchi, 1994; Kaufman et al., 2014; Kuang et al., 2015; Tanji & Evarts, 1976; Schaffelhofer et al., 2015; Gertz & Fiehler, 2015; Lindner et al., 2010; Verhagen et al., 2012).

In our experiment, participants had to execute one of two possible hand movements after a variable delay (open or close, see Fig. 1). During the delay, two aspects of the upcoming movement could be independently cued (or not): the movement type and the visuomotor mapping. We assumed that cueing (pre-specifying) movements would establish a specific motor plan during the delay; whereas in the uncued case, a motor plan would be initiated only ‘ad-hoc’ upon presentation of the execution prompt (Rosenbaum, 1980; cf. Ames et al., 2014; Pilacinski et al., 2018). Consequently, we expected relatively increased frontoparietal blood oxygenation level dependent signal (BOLD) during the delay period when participants were preparing for cued > uncued movements. If, alternatively, both movements were prepared in parallel during the delay, frontoparietal BOLD should show the opposite effect (uncued > cued).

**Figure 1.**
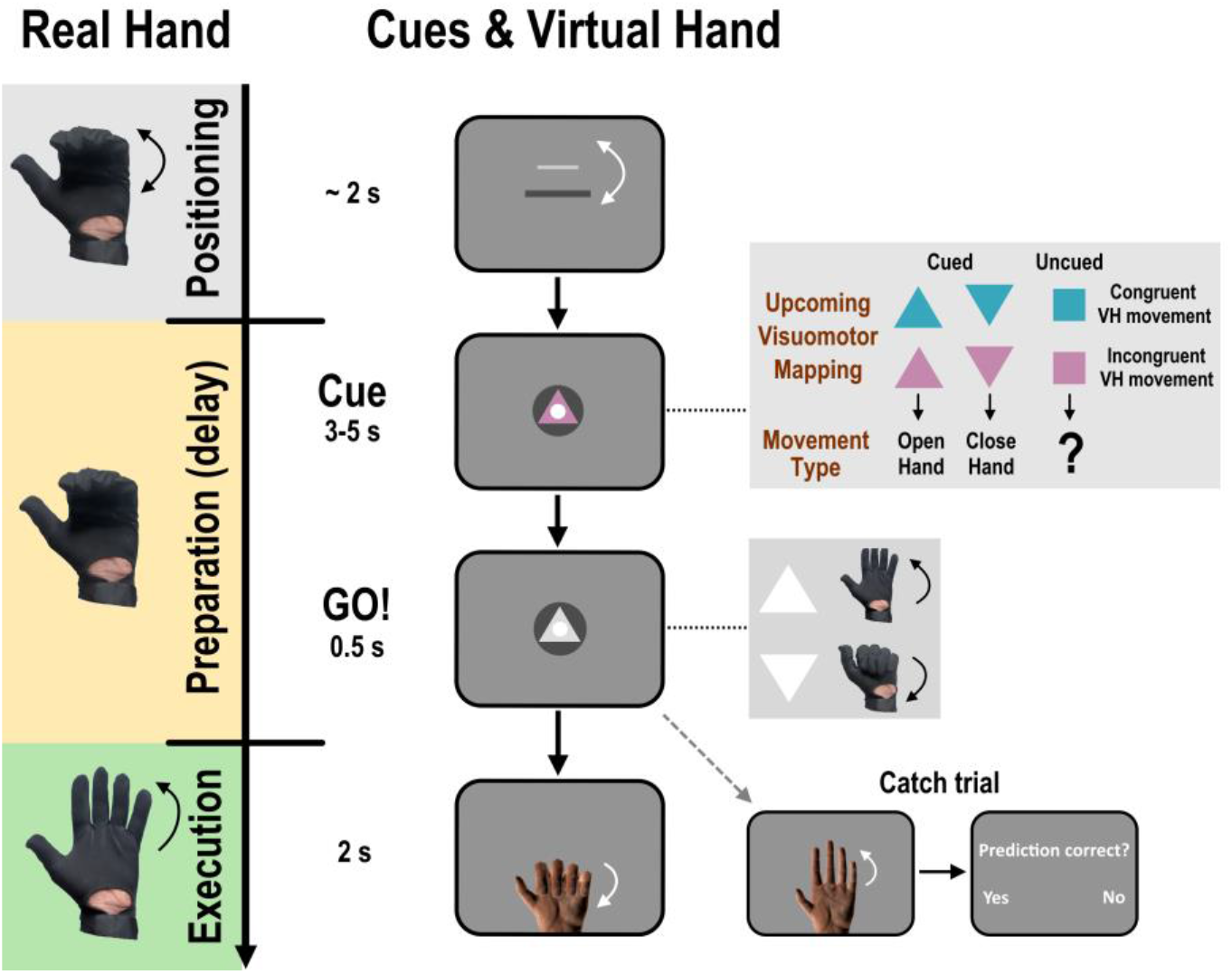
Experimental task to manipulate motor planning and visuomotor prediction. The participants’ real hand movements (left, hidden from view) were measured via a data glove and fed to a virtual hand (VH) model presented on screen (right). In each trial, participants had to execute one of two possible movements (open or close their hand) after a variable delay, starting from a neutral position (hand half-way closed). During the delay (preparation period), we presented a visual cue that could provide two kinds of predictive information: Firstly, the cue’s *shape* could specify the movement type to be executed after the delay (up- or downward triangle = open or close their hand, 25% of all trials each, i.e., together 50%) or leave it ambiguous (square = upcoming movement unknown, 50% of all trials). Secondly, the cue’s *color* indicated the to-be-expected visuomotor congruence; i.e., whether the virtual hand would next reflect the real hand movement (here: cyan cue) or the respective opposing movement (i.e., inverting the real hand movement, here: pink cue). Thus, we orthogonally manipulated processes related to motor planning and visuomotor prediction (the predicted mapping of visual movement feedback onto motor commands; i.e., what the hand movement will look like), respectively; in the absence of spatial movement targets during planning or execution. After a ‘Go signal’ and appearance of the virtual hand on screen, the participants executed the instructed hand movement while maintaining central fixation. In this particular example, the participant was cued that she would need to open her hand (triangle pointing upwards) and that the virtual hand would move incongruently during execution (pink cue), i.e., close. To ensure task engagement and attention to the cues, participants had to monitor and, in occasional catch trials, report whether the virtual hand movement matched the color-cued predictions (in 25% of trials, the virtual hand mapping was invalidly cued). Trials were separated by a brief (2s) interval. In this figure, the visual cues have been enlarged for display purposes.

By cueing incongruent (i.e., inverted) visual hand movements, we aimed to establish an anticipation of an unnatural visuomotor mapping. In other words, we assumed that participants would naturally expect visuomotor congruence, based on a life-long, learned association between executed movements and their visual feedback; and that incongruent movements would violate this expectation (Yon et al., 2018). When these ‘nonstandard’ visuomotor mappings were cued before the actual movement execution, we expected an anticipatory adjustment of the forward model’s predictions, detectable as increased delay period BOLD. Importantly, as our study design entailed no spatial targets during planning and execution, these predictions should specifically relate to visual body movement feedback, i.e.: *What my hand movement will look like*. Moreover, we expected motor planning to require more neuronal resources when integrating the (updated) predictions of nonstandard visual feedback in the state estimate—with corresponding interaction effects predominantly in the PPC and PMd.

## Materials and Methods

### Participants

30 healthy, right-handed volunteers (18 females, mean age = 25.4 years, range 19-35) participated in the experiment after signing informed consent. The experiment was approved by the ethics committee of the Technische Universität Dresden and conducted in accordance with this approval.

### Experimental design and procedure

Participants lay comfortably inside an MR scanner. On their left hand, they wore an MR-compatible data glove (5DT Data Glove 14 Ultra MRI, 60 Hz sampling rate, full-speed USB 1.1 connection), which measured each finger’s flexion via sewn-in optical sensors. The glove data were fed to a photo-realistic virtual hand model, which was thus controllable by the participants’ hand movements. We could not measure the exact intrinsic delay of the setup, but even including conservative estimates for USB transmission and projector latencies, the intrinsic delay should be less than visuomotor delays typically detected above chance level in such setups (e.g., 167 ms in our previous study using the same glove and hand model, Limanowski et al., 2017). The virtual hand received an average of the four sensor values to ensure smooth and coherent visual motion (cf. Limanowski & Friston, 2020). The participants’ real hand was placed outside of their view; i.e., during the entire experiment, participants only saw the virtual hand, but not their real hand. Prior to the experiment, the glove was calibrated carefully; i.e., the virtual hand model was shown and the participant moved her fingers to establish the possible range of flexion (minimum and maximum sensor values), and the virtual hand model’s finger flexion was adjusted to closely match the real hand posture and movement range according to the participant’s perception. If necessary, the calibration was repeated during the pauses between individual scanning runs. Through a mirror on the head coil, participants saw a screen, onto which a projector (Casio XJ-M156, Laser/LED hybrid, 1024 x 768 pixels resolution, 60 Hz refresh rate) displayed the virtual reality environment; i.e., the instructions, cues, and the virtual hand model. The virtual environment was created with the Blender 3D graphics software package (https://www.blender.org/, Version 2.79) and its Python programming interface. The experimenter observed the virtual display (via a mirrored screen) at all times, thus ensuring task compliance.

The participants’ task was to perform, in each trial, one of two types of cued hand movements (open or close, from a neutral starting position) after an instructed delay. During movement, the virtual hand model moved congruently or incongruently to the participants’ actual hand movements. Each trial consisted of two periods – a *preparation period,* in which participants could be cued about various aspects of the upcoming movement (i.e., inducing motor planning and visuomotor expectations, see below); followed by an *execution period*, where participants executed the hand movements, while receiving (condition-specific, see below) visual movement feedback from the virtual hand model. See Figure 1 for an example trial.

At the beginning of each trial, participants had to bring their real, unseen hand to the neutral starting position (i.e., fingers half-way closed). For this, a small white bar was presented on screen, whose vertical position was controlled by the participants’ hand posture via the data glove (i.e., the degree to which the hand was opened or closed). Participants had to align this bar with a central target (another bar, see Fig. 1); which, on average, took 1.96 s (SD = 0.48 s). Upon alignment, the bar was then replaced by a central white fixation dot for 1 s. Participants were instructed to keep their gaze on this fixation dot throughout the entire trial; i.e., until the end of the execution period.

Then, a cue was presented and remained visible throughout the delay period (3 to 5 s jittered, pseudo-randomized, evenly distributed across conditions). This cue could provide participants with two kinds of predictive information: The *type of the upcoming movement* (i.e., hand opening or closing) and the *congruence of the virtual and real hand movements* during this movement (see below). These two predictions were coded by cue shape and color, respectively. Thus, in half of all trials, the cue *shape* specified the movement that the participant would need to execute next. Specifically, in 25% of trials an upward-pointing triangle informed participants that they would have to *open* their hand, while in another 25% a downward-pointing triangle informed them they would have to *close* their hand. In the remaining half of all trials, the cue was square-shaped and, thus, did not reveal the to-be-executed movement type in advance. In other words, only in half of the trials participants were able to generate a specific motor prediction about the movement they would have to execute after the delay period.

Furthermore, the *color* of the cue was predictive of whether, during the upcoming movement, the virtual hand would move congruently (i.e., reflecting the participants’ real hand movement) or incongruently (performing the respective opposite, i.e., inverted, movement type) with the participant’s real hand movements. In the incongruent conditions, the values fed to the hand model from the glove’s sensors were inverted so that an opening movement would close the virtual hand, and vice versa. Thus, the movements of the virtual hand were controlled by the participant in both cases, ensuring matching virtual and real hand kinematics during congruent and incongruent conditions. This was important because our ‘incongruent’ virtual hand movements were, therefore, also predictable in principle from the real hand movements; i.e., from motor signals.

The two movement types were designed to produce, from the neutral starting position, a comparable range of visual motion; i.e., cover the same Euclidean distance on screen: At the beginning of the trial, the virtual hand model was displayed on the screen in the neutral starting position. In this position, the tip of the middle finger was positioned exactly in the middle of the screen (i.e., in the same position as the fixation dot); and the Euclidean distance between this fixation dot and the fingertip for either the fully opened or closed hand position was the same. While opening vs closing the hand were not diametrical opposites in terms of actual joint rotations (cf. Fig. 2A), all conditions contained the same number of open and close movements (see above); and all fMRI contrasts were calculated on conditional contrast estimates pooled over both movement types. Thus, kinetic or kinematic differences between opening vs closing movements were balanced across conditions; i.e., any potentially related effects were factored out in our fMRI analysis.

**Figure 2.**
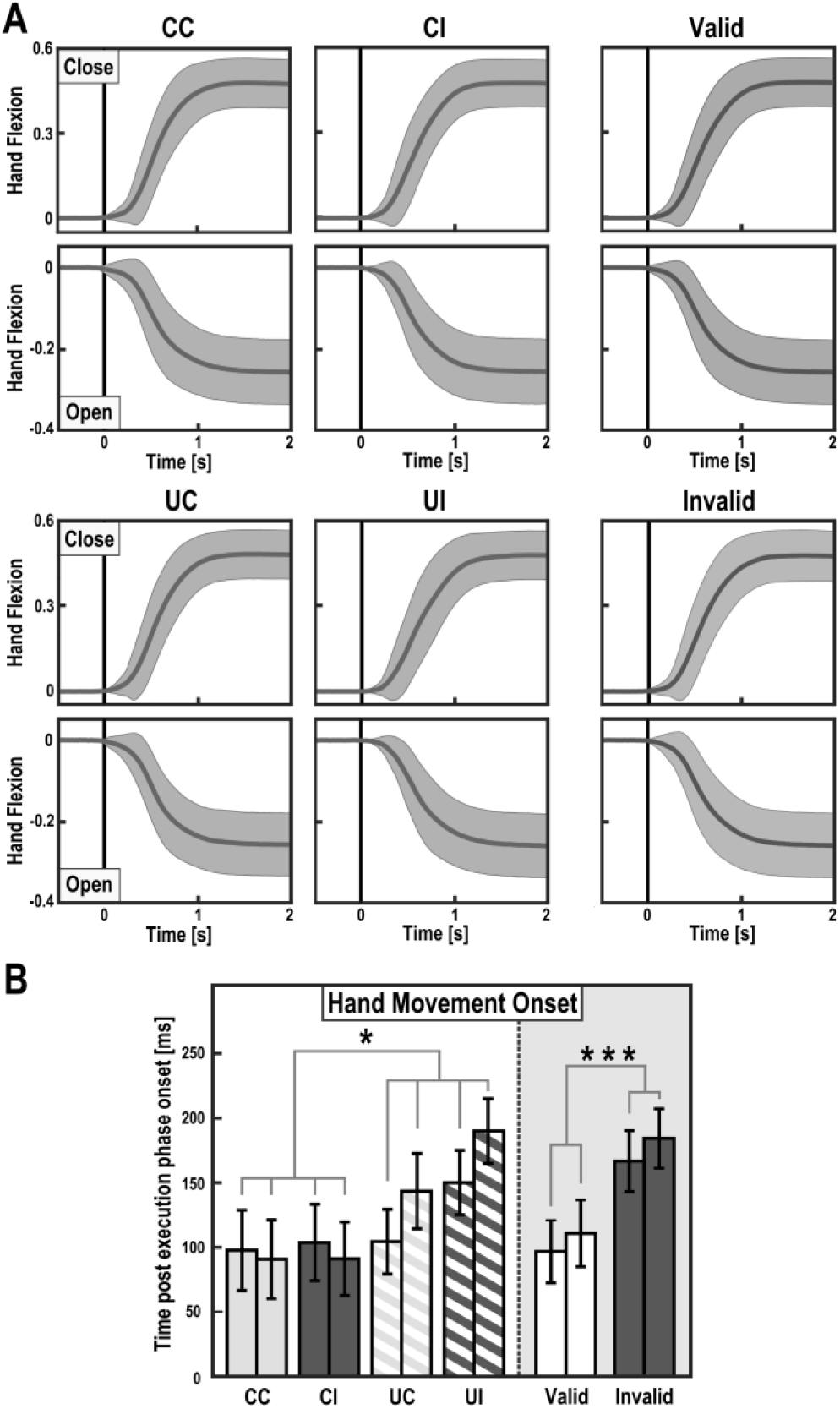
Average hand movements during the execution period. **(A)** The curves show the group-averaged unsmoothed flexion values with associated standard deviations as shaded areas (plotted for each condition, CC = cued/congruent, CI = cued/incongruent, UC = uncued/congruent, UI = uncued/incongruent, and separately for validly vs invalidly cued trials pooled over conditions). Positive flexion values indicating closing and negative values indicating opening, relative to the neutral starting position; time = 0 marks the instructed start and appearance of the virtual hand model. **(B)** The bar plots show the average movement onsets with associated standard errors of the mean (for each pair of bars, the first bar corresponds to the “open” hand movement, the second bar to the “close” movement). Movements were initiated significantly faster in cued > uncued trials, and in valid > invalid trials(\**p*<0.05, \*\*\**p*<0.001). See Results for details.

The cue colors (pink and cyan) were counter-balanced across participants. Following the preparatory period, a ‘Go’ signal (white triangle, centrally presented) appeared. Again, the pointing direction of this triangle instructed participants to open (triangle pointing up) or close (triangle pointing down) their hand. Note that, if the prior cue had not yet specified the movement type (i.e., square-shaped cue), the participants would only now know which movement they should execute.

Participants were instructed to execute the movement as soon as the ‘Go’ signal disappeared (after 500 ms) and, instead, the virtual hand model was presented on the screen. This was done to ensure that participants did not start to move reflexively before the virtual hand was visible. Participants then had to execute the hand movement fluently, for which they had up to 2 s (if no movement was made during that period, the trial ended). Participants were instructed and trained to make similar movements in all conditions and, after movement, to keep the hand in the final position (completely opened or closed) until the end of the trial. Then, the virtual hand disappeared again and a blank screen was shown for 2 s. As in the preparation period, participants still had to maintain fixation on the central dot during and post execution. In a prior training session, participants were able to practice the movements and familiarize themselves with all conditions. In sum, this experimental setup resulted in a factorial design with the factors *Movement type* (Cued, Uncued) and *Visuomotor mapping* (Congruent, Incongruent); and, correspondingly, four conditions: CC (cued movement with expected congruent mapping), CI (cued movement with expected incongruent mapping), UC (uncued movement with expected congruent mapping) and UI (uncued movement with expected incongruent mapping). In other words, movement planning was orthogonal to expectations about visuomotor congruence.

One important characteristic of our experimental design was the absence of spatial movement targets or ‘goals’ during planning or execution. The presence of such targets, during planning or executing movements towards (or away from) those targets, can make it difficult to distinguish activity related to the processing or encoding of the properties of visual movement feedback *per se* from, for instance, spatial eye-hand coordination or the visual-spatial representation of the movement goal (see Pilacinski et al., 2018; Lindner et al., 2010, for discussions). Therefore, in our design, we ensured that the visuomotor predictions pertained exclusively to the mapping of executed and observed hand movements, in other words: a prediction of what the planned hand movement would look like.

One implication of this design was that participants could have abstained from movement planning and simply waited for the ‘Go’ signal, presented in the form of a triangle. To ensure that participants nevertheless paid attention to the cues and, consequently, generated the respective visuomotor expectations or predictions, we included occasional trials in which the virtual hand moved *contrary* to the color-cued expectation (25% of all trials, balanced across conditions). For instance, the virtual hand moved congruently although the cue had predicted it to move incongruently (but note that the cues indicating the to be executed ‘own’ movement type were always valid). Following these invalidly cued trials, and following the same number of validly cued trials in each condition, participants had to report whether the virtual hand behavior matched the predictions of the cue color (cf. Yon et al., 2018). For this, a probe question (“Prediction correct?”) was presented centrally on the screen alongside two response options (“Yes”/”No”). Participants had to choose between those by pressing one of the response device buttons twice (2-Button Fiber Optic Response pad, Current Designs, placed in the participant’s right hand, on their lap). If they did so within 1.5 s, feedback (i.e. “right” or “wrong”) was shown for 0.75 s, otherwise they were visually instructed to respond faster the next time. A blank screen was shown for 2 s before the next trial. To increase motivation, participants received an additional monetary reward after the experiment, depending on the proportion of correct answers to the probe questions in the catch trials (1 € if at least 75% were answered correctly, 3 € for ≥85% and 5 € for ≥95%). We chose a relatively large number of catch trials (half of all trials in total) to maximize the task relevance of the cues during the preparation period; i.e., participants had to assume that the cue color could be probed by chance. Thus, the design for the execution period entailed *Visuomotor cue validity* (Valid, Invalid), as another nested factor (see below). This manipulation also allowed us to examine neuronal “surprise” responses to expectation violation; i.e., by comparing invalidly with validly cued catch trials.

Each trial lasted between 8.5 and 15 s. For each participant, we recorded 8 runs of 32 trials each (6.5- 8.5 mins run length), resulting in 256 trials (64 per condition and preparation/execution period, respectively) and around 60 mins scanning time in total (depending on hand positioning and response times). Participants were free to rest as much as they wanted in between the individual scanning runs.

### Behavioral data analysis

We tested for systematic differences in movement onset, duration, or amplitude between conditions; using the participants’ hand movement data from the glove sensors (averaged over all fingers but the thumb; the movements had a similar kinematic profile, comparable across conditions, without averaging the contribution of the individual fingers, see Results). We segmented the continuous glove data to select the execution period time window of each trial, smoothed (Gaussian-weighted moving average filter, time window 10 frames i.e. 167 ms), and re-adjusted the values to a baseline defined by the mean hand position during the 500 ms before appearance of the virtual hand (i.e. the instructed start of execution). Movement onset was defined, for each trial, as the respectively last time point before movement with zero velocity. Note that we could only approximate the actual movement onset from the recorded glove data; intrinsic delays due to the design of the glove (see above) should be taken into account when interpreting the onset values, but these delays were constant and hence irrelevant to our factorial analyses. The end of the movement was defined as the time point where hand velocity reached zero again. Movement duration and amplitude, respectively, were defined as the time difference and amplitude difference between movement onset and -end. Mean values for movement onset, duration and amplitude per condition were calculated for each subject. Based on these, a four-way ANOVA with the factors *Movement type* (Open, Close) *Movement type cueing* (Cued, Uncued), *Visuomotor mapping* (Congruent, Incongruent) and *Visuomotor cue validity* (Valid, Invalid) was used to test for differences between conditions. Reaction time in catch trials was defined as the difference between presentation of the question on screen and the time point at which participants confirmed their answer. All analyses were performed using MATLAB (MathWorks).

### FMRI data preprocessing and analysis

The fMRI data were recorded with a 3T scanner (MAGNETOM Prisma; Siemens) equipped with a 32- channel head coil. T2*-weighted images were acquired using an echo-planar imaging sequence (GRAPPA acceleration factor = 3, multi-band factor = 6, TR = 869 ms, TE = 38 ms, flip angle = 58°, voxel size = 2.4 mm^3^, matrix size = 88 x 88, 60 interleaved slices). Depending on the length of the individual session (see above), 3000-4000 functional image volumes were recorded per participant. Additionally, for each participant we recorded a T1-weighted anatomical image (3D, voxel size = 0.9 x 0.9 x 0.9 mm^3^, FOV = 272 x 272 mm, 240 slices, TR = 2.4 s, TE = 2.2 ms, flip angle = 8°) and two GRE field maps (64 layers, voxel size 2 x 2 x 2 mm^3^; matrix size = 88 x 88; TR = 698 ms, TE1 = 5.19 ms, TE2 = 7.65 ms, flip angle 54°), after the second and sixth scanning run, respectively. All fMRI preprocessing steps and analyses were performed using MATLAB (MathWorks) and SPM12 (Wellcome Trust Centre for Neuroimaging, University College London; https://www.fil.ion.ucl.ac.uk/spm). The functional images were realigned and unwarped using voxel displacement maps calculated from the two field maps (the first map was used for runs 1 to 4, the second for runs 5 to 8). These images were then co-registered with the individual T1 image, spatially normalized to MNI space via previously estimated deformation fields, resliced to 2 mm³ voxels, and smoothed with an 8 mm full width at half maximum Gaussian kernel. The T1 images were also normalized (but not smoothed) and averaged to create a group averaged structural image.

In our fMRI analysis, we adopted a mass univariate approach; i.e., we tested for voxel-wise differences in the amplitude of the BOLD signal between our experimental conditions. Implicit in this is the assumption that a relatively higher or lower BOLD signal in one condition compared to another (i.e., a differential contrast) indicates a regionally in- or decreased metabolic demand in this condition, respectively. We interpret such differences as evincing an increased engagement of brain regions in experimentally manipulated processes related to motor planning and visuomotor prediction, as identified per our factorial between-condition comparisons (see Fernandez-Ruiz et al., 2007; Lindner et al., 2010; Bernier et al., 2012; Gertz & Fiehler, 2015; or Pilacinski et al., 2018, for similar approaches).

For each participant, we fitted a general linear model (GLM, 128 s high-pass filter, including temporal and dispersion derivatives for each regressor) to the preprocessed data. We included stick (0 s duration) regressors modeling each combination of the two factors *Movement type* (Cued, Uncued) and *Visuomotor mapping* (Congruent, Incongruent) in the preparation and movement period, respectively. The resulting four regressors were thereby aligned with the onsets of the predictive cues; or with the onsets of the movement execution period including the virtual hand presentation. To model any potential ‘surprise’ effects, the regressors of the execution period were each supplemented by a mean-centered parametric modulator encoding trials in which the virtual hand moved unlike cued with +1, and trials in which it moved like cued with -1. Thereby, we applied the modulation to the catch trials only; i.e., ensuring the surprise effect was calculated on the same amount of trials (all with subsequent button presses) in invalidly and validly cued movements.

We added the six realignment parameters as noise regressors to each run, alongside a session constant. Furthermore, we added three regressors of no interest modelling the differences in movement amplitude, movement duration, and reaction time (i.e., the time it took participants to initiate a movement after onset of the execution period) during the execution period. Each kinematic regressor modelled trial-to-trial variations (in amplitude, duration, and reaction time) across the entire scanning session; i.e., it also modelled differences between conditions. Two additional regressors of no interest were included: one modeling the initial hand positioning (a 0s duration stick regressor with an additional orthogonalized parametric regressor encoding the duration from trial start until the correct hand position was reached and the fixation dot presented), and one modeling the response periods after the occasional catch trials (a 0s duration stick regressor with an additional orthogonalized parametric regressor encoding the respective variations in reaction time until the confirmatory key press). On the first (single participant) level, T-contrast images were created for each regressor vs baseline, across all 8 runs per participant (i.e., over all 64 trials per preparation or execution period per condition. These first-level contrast images were entered into group-level GLMs using a flexible factorial design, in which the group (second-level) T- and F-contrasts were calculated. We thereby used 2x2 factorial designs with the aforementioned factors to evaluate the main effects of cued vs uncued movements and congruent vs incongruent visuomotor mapping, as well as their potential interaction, during the preparation and execution periods, respectively.

We furthermore tested for the *consistency* of main effects; i.e., whether a given main effect of one experimental factor was significant at each level of the respective other factor. To do this, we calculated ‘null’ conjunction contrasts; a statistical test whether a BOLD signal difference in a given voxel is significant in each of multiple tested contrasts. Thus, we calculated a conjunction across the contrasts CC>UC and CI>UI (testing for consistent significance of the main effect movement type cued > uncued across both levels of the factor visuomotor mapping); and another one across the contrasts CI>CC and UI>UC (testing for consistent significance of the main effect incongruent > congruent visuomotor mapping across both levels of the factor movement type cueing). These conjunction contrasts were evaluated for statistical significance using a voxel-wise threshold of *pFWE*<0.05, corrected within the restricted search region defined by the significant voxels obtained from the respective main effect.

Furthermore, we tested for any influence of the (cued) preparatory processes on activity in the respectively identified areas during subsequent movement execution. As we did not have any specific prediction about the directionality of response differences during movement execution, we used non-directional F-contrasts, with post-hoc t-tests to examine directionality, in this case. To assess potential surprise effects during execution associated with invalidly > validly cued movements, we calculated an analogous 2x2 factorial group level design on the parametric ‘surprise’ modulators of each condition (i.e., on T contrast images weighting invalidly cued catch trials in each condition with +1 and validly cued ones with -1, see above). Finally, supplementing our main analysis (which focused on differences in metabolic demand via estimating the amplitude of the hemodynamic response), we looked for analogous conditional differences in the loadings on the temporal and dispersion derivatives of our model’s regressors; using an analogous second-level factorial design as in the main (amplitude) analysis on the first-level contrast images of the respective derivative. These estimates can provide additional information about the shape of the BOLD signal; i.e., a relatively larger estimate of the temporal derivative may indicate a relatively earlier hemodynamic response, and a relatively larger estimate of the dispersion derivative may indicate a relatively narrower response (Friston et al., 1998; Henson et al., 2002). This can supplement the interpretation of the hemodynamic (amplitude) response in terms of underlying neuronal, e.g. predictive, processes (cf. Kavroulakis et al., 2022).

Activations in the whole brain obtained from the group-level contrasts were assessed for statistical significance using a voxel-wise threshold of *p*<0.05, family-wise error corrected (*p*FWE<0.05). In some cases, we had specific regions of interest defined a priori by orthogonal contrasts, and restricted the search space accordingly, thus applying a small volume correction under the same voxel-wise threshold of *p*FWE<0.05 for those results (similar to the conjunction contrasts described above): Firstly, we looked for an interaction of motor and visuomotor planning in brain regions showing a response to both individual effects. Therefore, we defined the search space for these interaction effects based on all (whole-brain) significant voxels obtained from the null conjunction of the respective main effects (Fig. 5). For completeness, we also tested for whole-brain results. Based on the previously reported overlap between action planning and execution (Lotze et al., 1999; Ames et al., 2014), we expected that brain regions involved in motor or visuomotor planning would also show condition specific effects during movement execution. Therefore, for the analysis of responses during the execution period, we restricted the search space for effects to all significant voxels obtained from each respective analogous contrast during the preparation period. Thirdly, we hypothesized that surprise effects might emerge in brain regions implementing the forward model, i.e., issuing the respective forward predictions. In other words, in those regions, we expected a relative increase in BOLD in response to a (sensory) prediction error elicited by the unpredicted visuomotor mapping. As the generation of those predictions should be highlighted by the main effect of preparation for incongruent > congruent movements (see Introduction), we used those significant activations as a mask image to look for corresponding surprise effects; but also calculated an additional whole-brain analysis.

The significant activation differences resulting from the group level contrasts are presented as renders on SPM12’s template brains, or superimposed onto the group averaged normalized structural image. The displayed significant activations contain only voxels surviving the chosen threshold of *p*FWE<0.05.

The contrast estimate plots were created by extracting and averaging the first-level contrast estimates for each condition from the peak voxel of each anatomically distinct cluster, using the rfxplot toolbox (Gläscher, 2009). For anatomical reference, we used the SPM Anatomy toolbox (Eickhoff et al., 2005).

### Eye tracking control experiment

Our non-spatial (i.e., without spatial movement targets) movement task was intended to render eye movements useless and, thus, unlikely. As we did not measure gaze on-line inside the scanner, we conducted an eye-tracking control experiment post-hoc, with the same design. The aim of this experiment was to determine whether participants could, in principle in this task, maintain fixation onto the central cues/dot; and whether gaze behavior was comparable across experimental conditions. A sub-sample of our fMRI study participants (*N*=6) completed the same task as in the scanner (with 4 instead of 8 runs in total), with stimuli presented at a size corresponding to those seen in the scanner environment. Participants were seated comfortably in a constantly lit room in front of a screen (at a distance of 93 cm) with their head stabilized via a chin rest, while we measured gaze position with an Eyetracker (EyeLink 1000 Plus, Desktop Mount, SR Research). We segmented the resulting eye position data to select 2500 ms time windows for both the preparation and execution periods of each trial, with phases of data loss (e.g. due to eye blinks) corrected via linear interpolation. Gaze position was defined relative to the position of the central fixation dot; i.e., the measured fixation position in the horizontal and vertical planes was combined to a measure of Euclidean distance from the fixation point, and converted to degrees of visual angle. To evaluate the quality of fixation, we tested for differences in mean gaze position and in the variability of fixation (i.e., the standard deviation); and we tested for this separately for the two possible movements. Thus, we performed a four-way repeated-measures ANOVA with the factors *Movement* (Open, Close), *Movement type* (Cued, Uncued), *Visuomotor mapping* (Congruent, Incongruent), and *Visuomotor cue validity* (Valid, Invalid) on the participants’ condition averages of both fixation position and variability, respectively.

### FMRI right-hand control experiment

To follow up on the finding that the main effects of cueing (and to a lesser extent also execution) were more pronounced in the left than in the right PPC (i.e., ipsilateral to the moving hand, see Figs. 3-6), we scanned 6 of our original participants again in a separate session; using an identical experimental setup and design (with 5 instead of 8 runs in total) but with right-hand instead of left-hand movements (and a right instead of left virtual hand model). These data were preprocessed and analyzed as described for the main experiment. As the small sample size limited the power of the random-effects group analysis, we evaluated and displayed the resulting statistical parametric maps at a threshold of *p* < 0.001, uncorrected (see supplement, Fig. S6).

**Figure 3.**
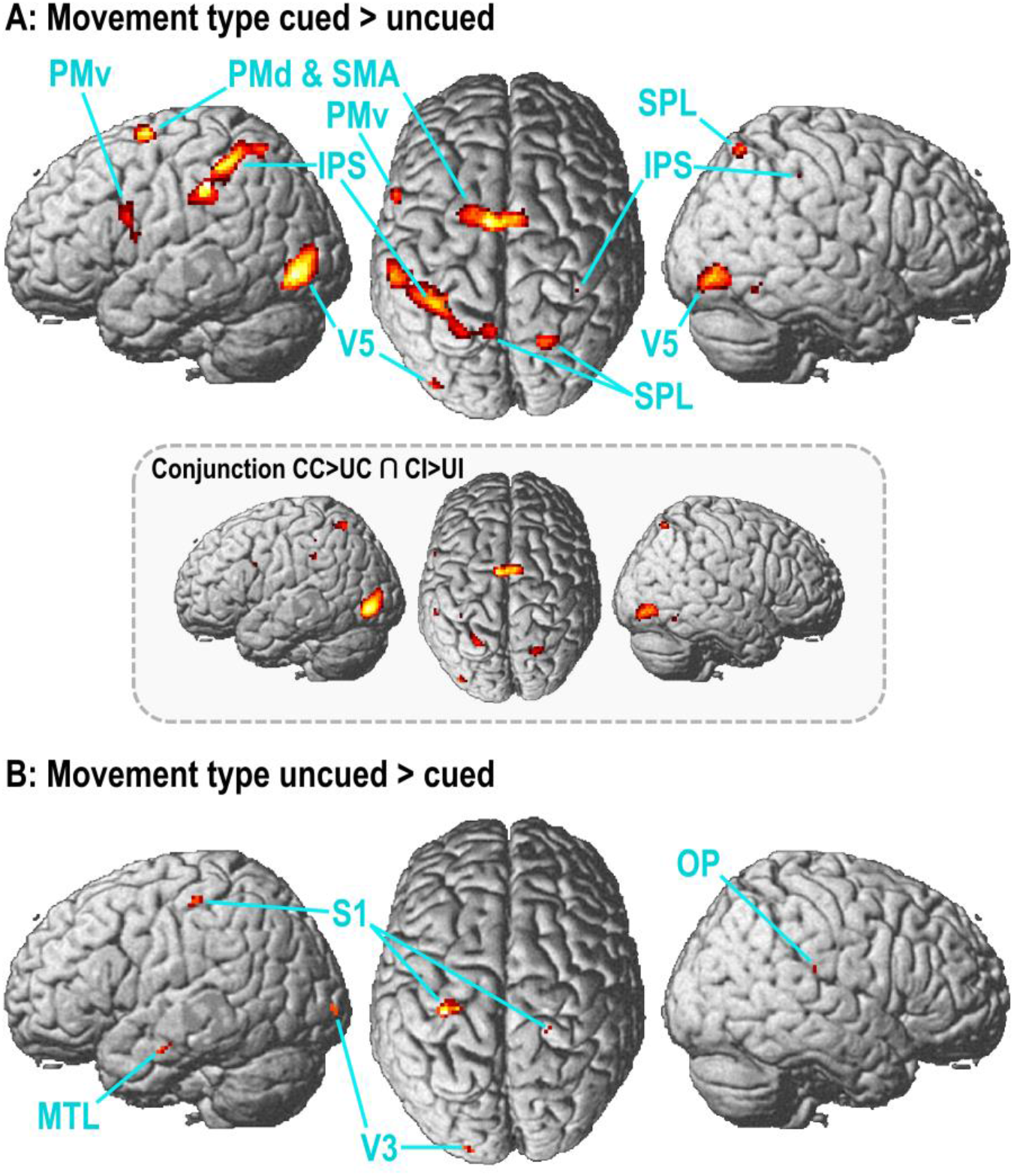
Brain activations related to motor planning during the delay period. **(A)** The renders show significant (*p*FWE<0.05) voxels obtained from contrasting the delay period of trials in which the to-be-executed movement type was cued, with trials in which this was left ambiguous until execution (cued > uncued). Inlay: Significant (*p*FWE<0.05) voxels obtained from a conjunction testing for consistency of the effect cued > uncued across congruent and incongruent mappings. **(B)** The converse contrast (uncued > cued) yielded significant in visual and somatosensory cortices. CC = cued/congruent, CI = cued/incongruent, UC = uncued/congruent, UI = uncued/incongruent. See Table 1 for details.

**Figure 4.**
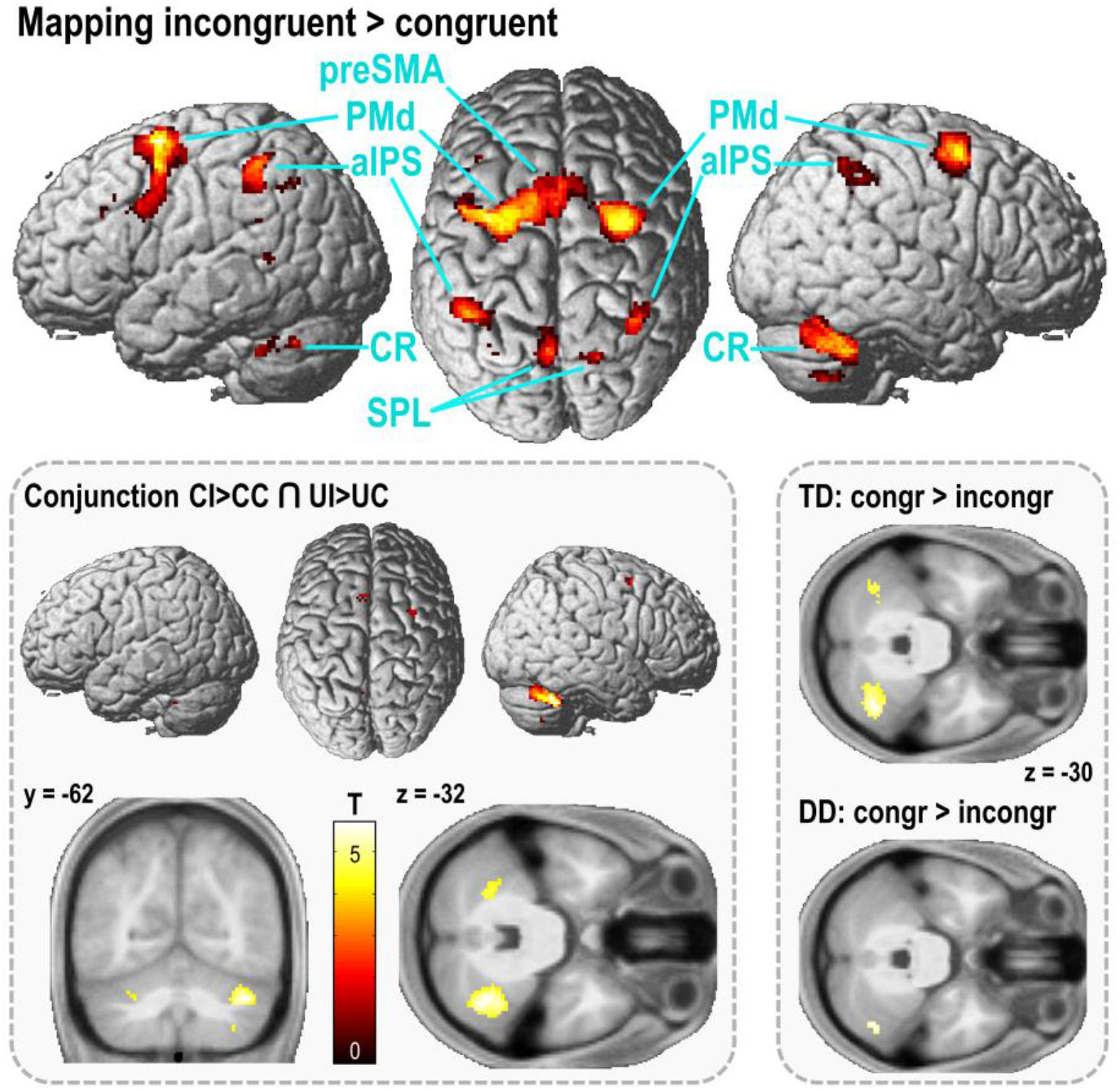
Brain activations related to visuomotor prediction during the delay period. The renders show significant (*p*FWE<0.05) voxels obtained from contrasting the delay period of trials in which the cue color predicted incongruent, compared with congruent visuomotor mapping (incongruent > congruent). The converse contrast yielded no significant effects. Inlay, left: Significant (*p*FWE<0.05) voxels obtained from a conjunction testing for consistency of the effect incongruent > congruent visuomotor mapping across cued and uncued movements. CC = cued/congruent, CI = cued/incongruent, UC = uncued/congruent, UI = uncued/incongruent. CR = cerebellum. See Table 1 for details. Inlay, right: An analysis of the loadings on the temporal derivative (TD, top) and dispersion derivative (DD, bottom) revealed voxels in corresponding areas of the cerebellum showing significantly later and wider hemodynamic responses when incongruent movements were prepared.

**Figure 5.**
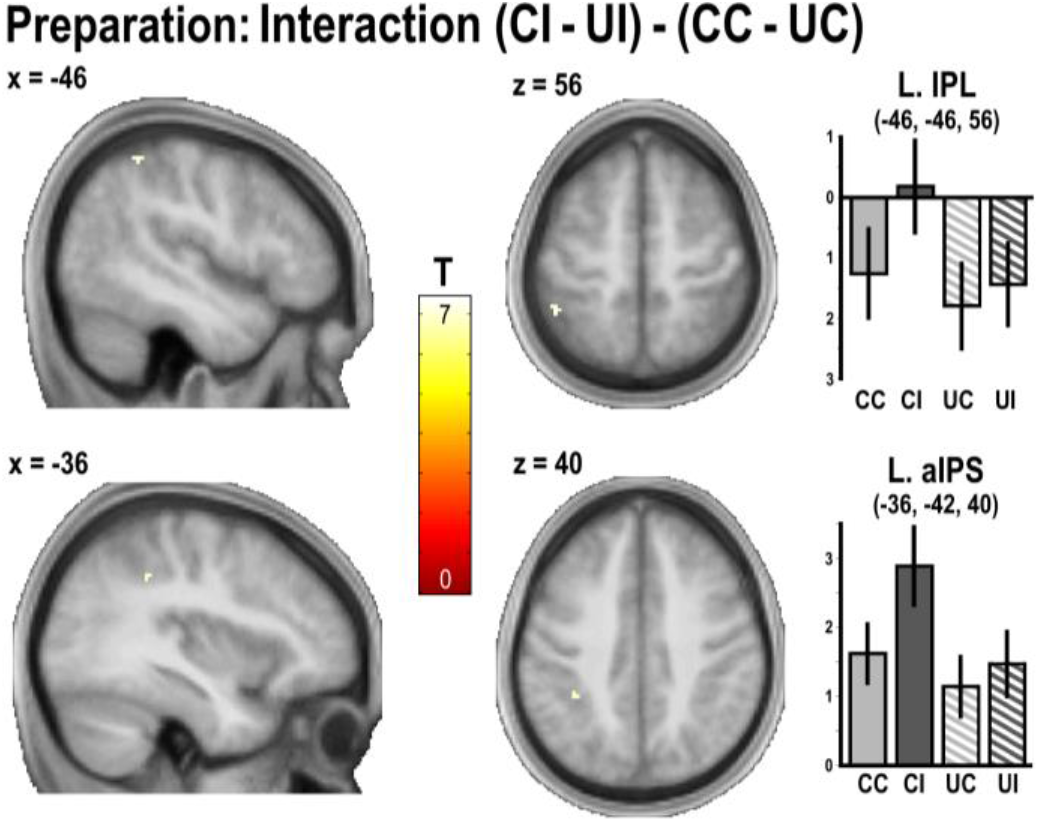
During the delay period, voxels in the left IPS/IPL showed a significant (*p*FWE<0.05) interaction effect, within the areas showing both main effects as identified by a null conjunction contrast (see main text): the highest responses were observed during the preparation of specified (cued) movements when an incongruent visuomotor mapping was expected. The bar plots show the contrast estimates with associated standard errors of the mean. CC = cued/congruent, CI = cued/incongruent, UC = uncued/congruent, UI = uncued/incongruent.

**Figure 6.**
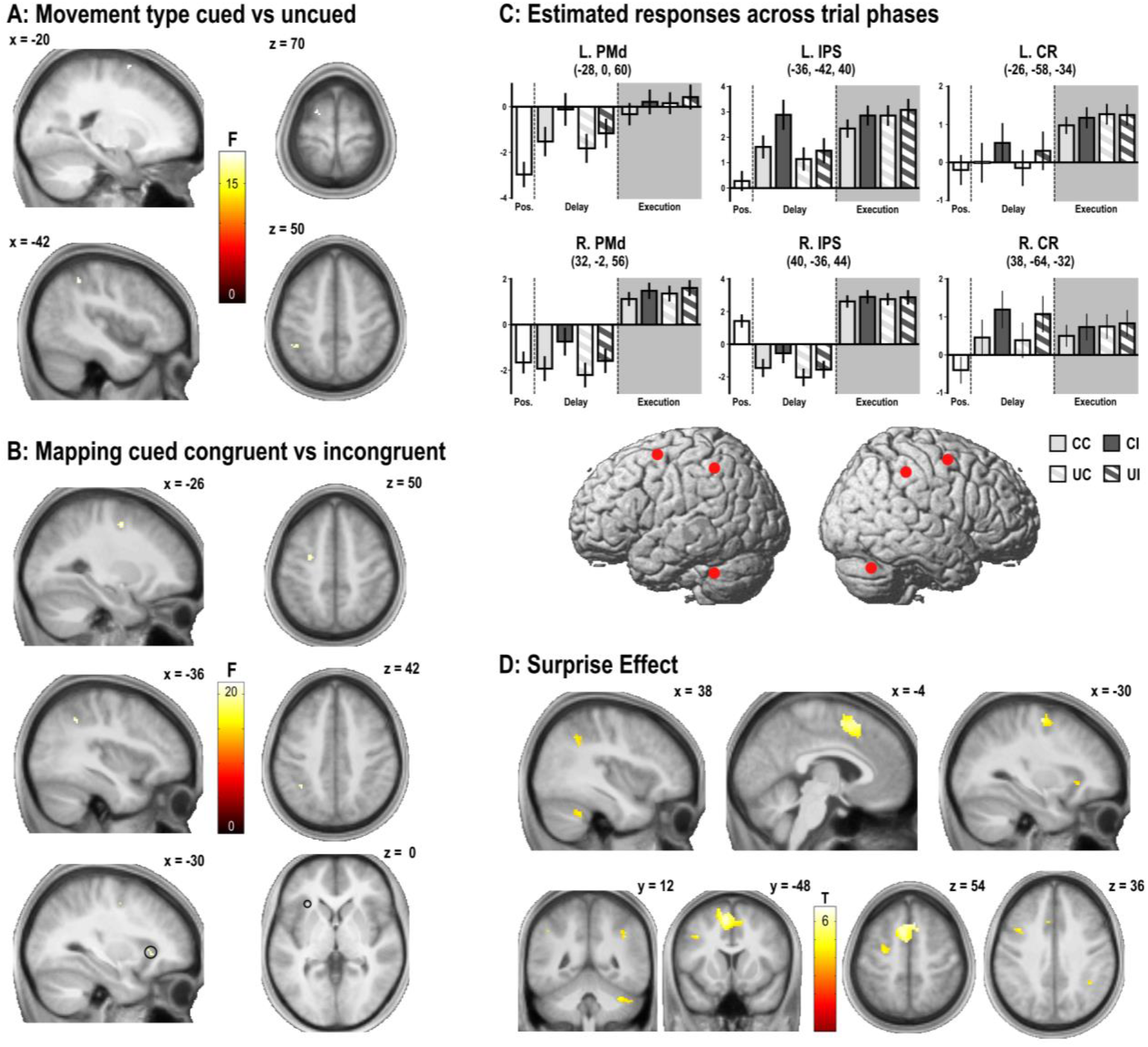
Brain activation differences during movement execution, related to prior movement planning and visuomotor predictions. **(A)** Voxels showing significant BOLD signal differences during movement execution in trials where the movement type had been cued in advance versus when it was left ambiguous (F-test, cued vs uncued, *p*FWE<0.05; a post-hoc t-test revealed significantly stronger responses for uncued > cued, see Results). **(B)** Voxels showing significant BOLD signal differences during movement execution in trials where congruent vs incongruent visuomotor mapping had been cued (F-test, congruent vs incongruent, *p*FWE<0.05; a post-hoc t-test revealed significantly stronger responses for incongruent > congruent, see Results). **(C)** Estimated hemodynamic responses, throughout the different trial phases of the experiment, in the main frontoparietal and cerebellar areas identified in our analysis (the renders show the peak coordinates marked with red dots). The plots show contrast estimates with associated standard errors of the mean: The first bar shows the estimate for the pre-positioning period prior to each trial (cf. Fig. 1); the next bars show the contrast estimates for each condition, first for the delay (preparation period), then for the execution period. CC = cued/congruent, CI = cued/incongruent, UC = uncued/congruent, UI = uncued/incongruent. **(D)** Significantly (*p*FWE<0.05) stronger activations by unexpected visuomotor mappings (‘surprise’ effect) within brain areas showing effects of visuomotor prediction (cf. Fig. 3B and Fig. S5). CC = cued/congruent, CI = cued/incongruent, UC = uncued/congruent, UI = uncued/incongruent. See Table 3 for details and supplementary Figures S4-S5 for (uncorrected) whole-brain results.

**Table 1.**
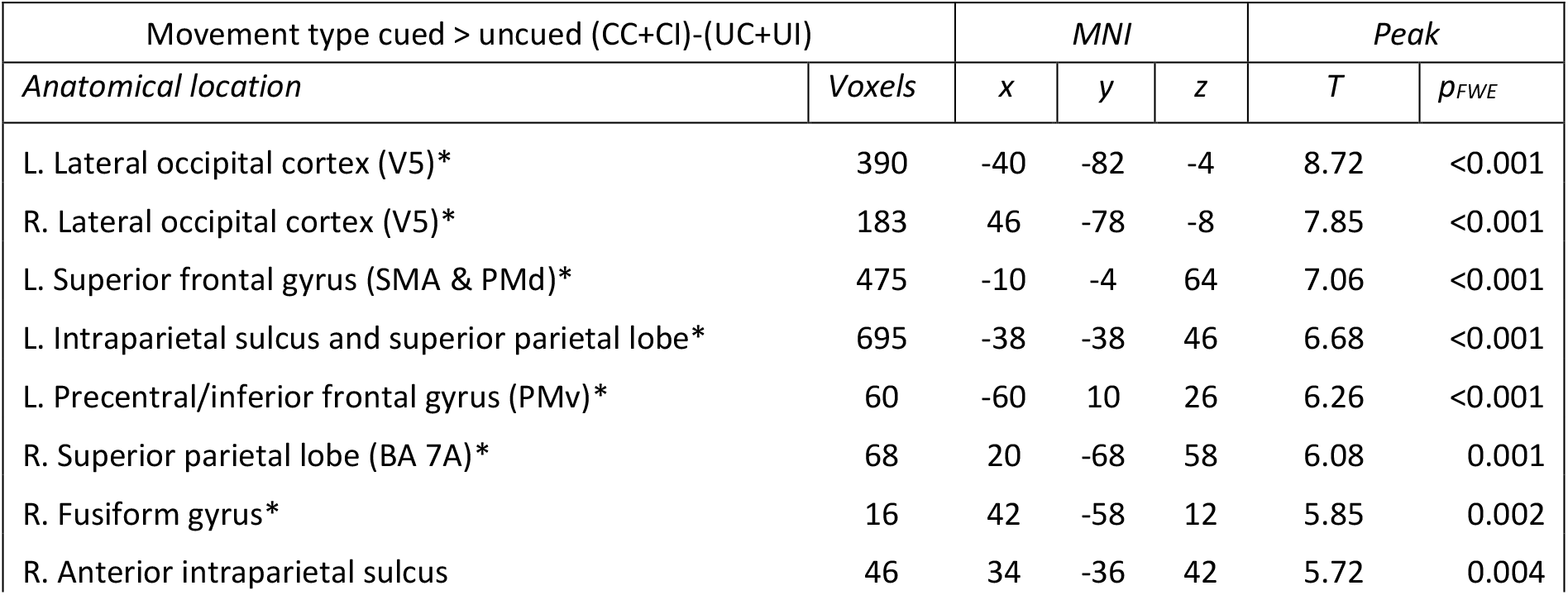
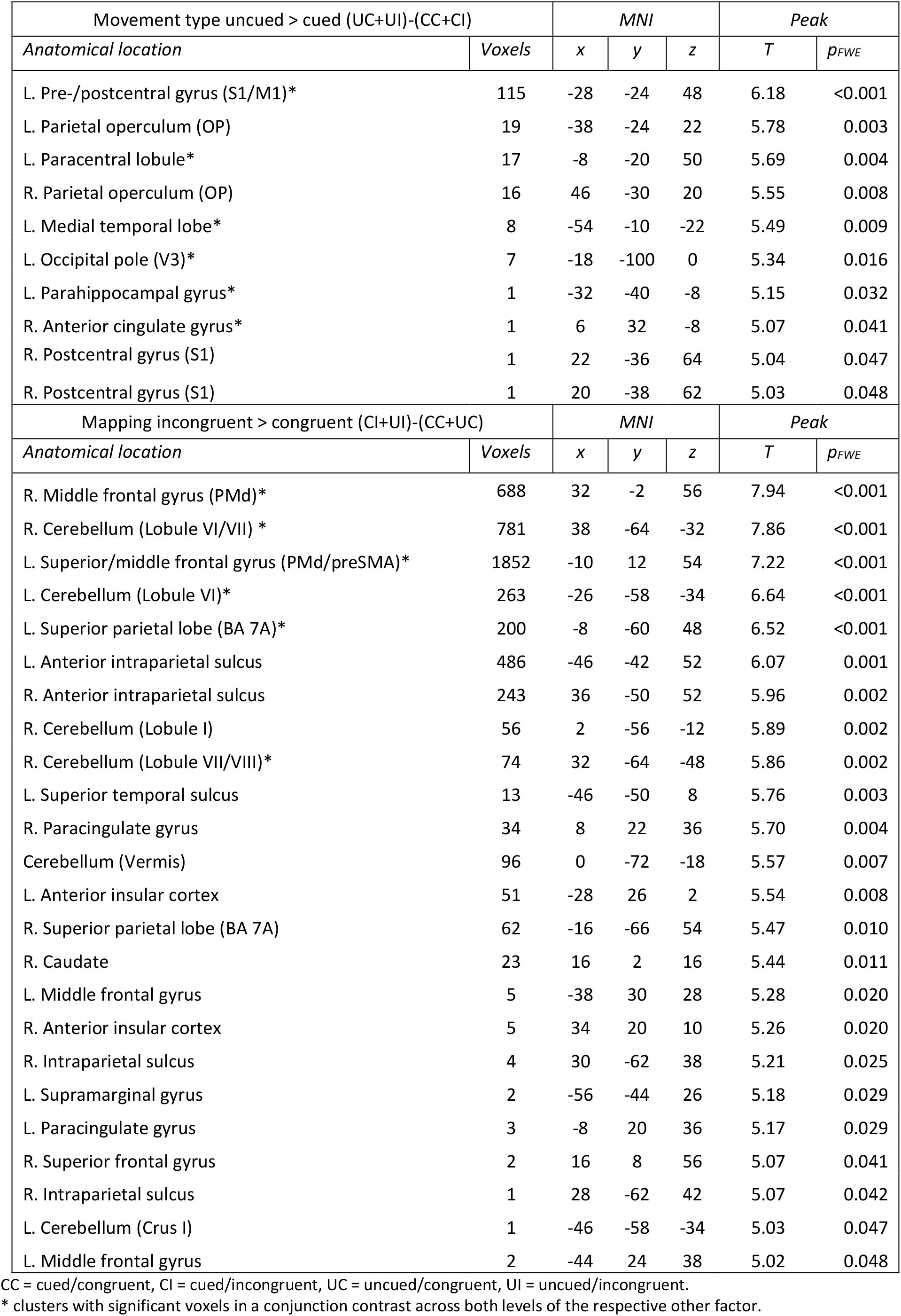
Brain areas showing significant main effects during the delay.

## Results

All participants showed very good task performance; i.e., all performing >97% correct movements (on average, only one incorrect movement per recording session) and correctly identifying >93% of the catch trials (on average, >98% of responses were correct). The average reaction time in catch trials was 0.920±0.246 s. There were no significant effects of visuomotor cue validity, visuomotor mapping, or motor planning on reaction times. The participants’ hand movements were consistent across conditions (Fig. 2). On average, movement execution was detectable from the glove data 121±119 ms after the onset of the execution period, with a mean movement duration of 1181±316 ms. Movement amplitude, in terms of average flexion values relative to the starting position, averaged to -0.26±0.08 (hand opening) and 0.48±0.09 (hand closing), respectively. Open vs close hand movements differed in terms of movement amplitude (F(2,29) = 73.880, *p*<0.001), but not in terms of onset or duration of the movements. Crucially, there was no significant effect of visuomotor mapping (congruent vs incongruent) on movement onset, duration, or amplitude of the hand movements. Likewise, motor planning (cued vs uncued) had no significant effect on movement amplitude or duration. However, movements in cued trials were on average initiated 51 ms earlier than those in uncued trials (F(2,29) = 7.34, *p*=0.011). This very likely indicates a reaction time benefit of prior motor planning in cued trials (which speeded up subsequent execution upon prompting), in line with previous findings (Rosenbaum, 1980; cf. Ames et al., 2014). Finally, there was a significant three-way-interaction (cueing x mapping x cue validity) on movement duration (F(2,29) = 6.82, *p*=0.014), which was, however, difficult to interpret. Interestingly, movements in invalidly cued visuomotor mapping trials were, on average, initiated 72 ms later (F(2,29) = 42.09, *p*<0.001) and 155 ms shorter (F(2,29) = 56.67, *p*<0.001) than movements in validly cued trials; but movement amplitude was unaffected by cue validity (F(2,29) = 0.38, n.s). This could indicate very early surprise responses to the expectation violation, leading to a slower onset (which was defined as the glove sensor flexion data crossing a certain threshold, see Materials and Methods). Importantly, however, this effect was comparable across experimental factors (i.e., no significant interaction with either visuomotor mapping or motor planning), hence our fMRI contrasts of interest – between conditions – were unlikely to have been affected by this. Furthermore, kinematic differences between trials and conditions were included as covariates of no interest in the fMRI analysis (see Materials and Methods). All of the above significant conditional effects were replicated when analyzing the data of the four fingers individually (significant for the index, middle, and ring finger, *F*s > 5.9, all *p*s < 0.05; non-significant for the pinky, *F*s > 1.9, *p*s < 0.17); there were no additional significant main or interaction effects when looking at each finger individually. A supplementary eye-tracking experiment (*N*=6) showed that participants were, in principle, able to maintain very stable fixation onto the central cues independently of condition throughout the preparation period, and overall also during movement execution (see Fig. S1 for details).

In sum, the participants’ behavior suggested a high task engagement and comparable hand movements (and fixation) across conditions, while showing a likely reaction time benefit of pre-cueing movements. Moreover, the almost perfect responses to catch trials, in which the validity of the color-cued and the actual visuomotor mapping was probed, indicated that participants paid attention to the visuomotor cues. Thus, we were confident that participants generated the desired (cued) motor plans and visuomotor predictions during scanning.

In our fMRI analysis, our primary aim was to identify brain areas involved in motor planning and visuomotor prediction; i.e., we looked for corresponding BOLD signal differences during the delay period. First, we contrasted the delay period of trials in which a specific movement type could be prepared – based on the cue shape – with trials in which this was not possible (i.e., main effect of movement type cued > uncued). As expected, this contrast revealed significantly (*p*FWE<0.05) stronger BOLD signal predominantly in the bilateral frontoparietal cortices; including the bilateral aIPS (and, in the left hemisphere, larger portions of the IPS), the bilateral SPL, the SMA, medial parts of the bilateral PMd, the left ventral premotor cortex (PMv); and the bilateral lateral occipital cortices (LOC, including area V5). See Table 1 and Figure 3A for details. A conjunction analysis revealed that, in almost all of these regions, the main effect of cued > uncued movements was consistent across congruent and incongruent visuomotor mappings (i.e., levels of the second experimental factor; *p*FWE<0.05, see Table 1). In other words, this suggested a consistently elevated BOLD signal by motor planning during preparation for congruent and incongruent visuomotor mappings alike. None of these regions showed significant differences in an analysis of the analogous contrasts calculated on the temporal and dispersion derivatives (Fig. S2). The converse contrast (i.e., delay period of uncued > cued movement types) revealed smaller clusters of significant voxels mainly in the visual and somatosensory cortices (Table 1 and Fig. 3B).

Then, we tested for increased responses during the delay period of trials in which an incongruent > congruent mapping of the virtual hand movements relative to the real hand movements was cued; assuming that here, predictions of the forward model would need updating. There was a significant main effect of cues predicting an incongruent > congruent mapping in the cerebellum (centered on Lobule VI in each hemisphere), premotor (the bilateral PMd, and the left preSMA and PMv) and posterior parietal (the left aIPS and the bilateral SPL) areas (see Figure 4 and Table 1 for all effects). However, a conjunction analysis revealed that this effect was consistent across cued and uncued movement types in only some of those regions (*p*FWE<0.05, see Fig. 4): Most notably, the strongest consistent effects across the CI > CC and the UI > UC contrasts were located in the bilateral Lobules VI of the cerebellum (with the strongest effect located in the right cerebellum, see slice overlays in Fig. 4). Only few other voxels in the PMd, the SMA, and in the left precuneus showed consistent effects. No significant activation differences were observed for the converse contrast, cued congruent > incongruent visuomotor mapping. Furthermore, an analysis of the analogous contrasts calculated on the temporal and dispersion derivatives revealed significantly earlier and narrower hemodynamic responses related to the preparation of congruent > incongruent mappings in the same areas of the cerebellum (Fig. 4; further temporal but not dispersion effects were observed in the left IPS, PMd, and PMv, see Fig. S2). In other words, the (right) cerebellum’s hemodynamic response peaked relatively later and was wider during the preparation for visuomotor incongruence, compared with congruence—suggesting, together with its higher amplitude, more sustained neural activity (cf. Friston et al., 1998; Henson et al., 2002).

Next, we specifically tested for brain areas that would jointly encode both preparatory processes in our design; i.e., those related to motor planning and visuomotor mappings. A null conjunction across both main effects (cued > uncued movement type and incongruent > congruent visuomotor mapping, Fig. 3A and Fig. 4) revealed consistently significant voxels within the bilateral aIPS and SPL, and the left medial PMd/SMA (*p*FWE<0.05, Table 2 and Fig. S3). Then, we evaluated our hypothesis that motor planning and visuomotor prediction would interact; i.e., we tested for relatively stronger movement planning effects (cued > uncued movements) under expected visuomotor incongruence, compared with congruence; i.e., via the contrast (CI-UI)-(CC-UC). Only the left aIPS/IPL showed a significant corresponding interaction effect: a relatively stronger increase in fMRI activity for delay periods in which the cue predicted a specific movement type with incongruent > congruent visuomotor mappings (CI>CC), compared with when the movement type was left ambiguous (UI>UC; see Figure 5 and Table 2). In other words, in the left aIPS, while the main effect of cueing itself was consistent, the main effect of preparing for incongruent visuomotor mappings was significantly stronger in the cued movement conditions. No other significant effects were found; a similar interaction effect in the left PMd did not reach significance (Fig. S3).

**Table 2.**
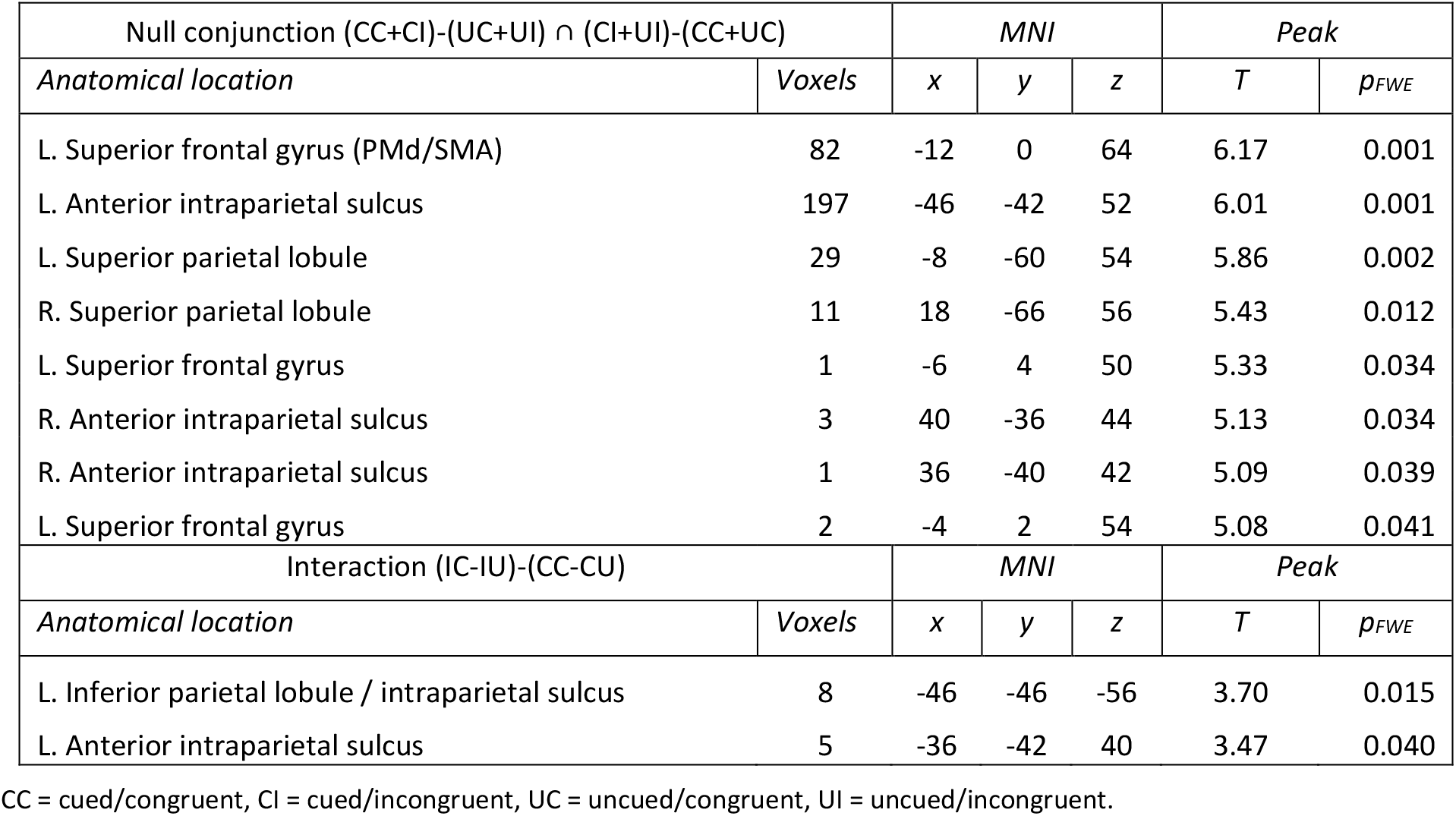
Significant activation differences obtained from the conjunction of main effects and the interaction effect.

| Null conjunction (CC+CI)-(UC+UI) ∩ (CI+UI)-(CC+UC) | | MNI | | | Peak | |
| --- | --- | --- | --- | --- | --- | --- |
| Anatomical location | Voxels | x | y | z | T | $p_{FWE}$ |
| L. Superior frontal gyrus (PMd/SMA) | 82 | -12 | 0 | 64 | 6.17 | 0.001 |
| L. Anterior intraparietal sulcus | 197 | -46 | -42 | 52 | 6.01 | 0.001 |
| L. Superior parietal lobule | 29 | -8 | -60 | 54 | 5.86 | 0.002 |
| R. Superior parietal lobule | 11 | 18 | -66 | 56 | 5.43 | 0.012 |
| L. Superior frontal gyrus | 1 | -6 | 4 | 50 | 5.33 | 0.034 |
| R. Anterior intraparietal sulcus | 3 | 40 | -36 | 44 | 5.13 | 0.034 |
| R. Anterior intraparietal sulcus | 1 | 36 | -40 | 42 | 5.09 | 0.039 |
| L. Superior frontal gyrus | 2 | -4 | 2 | 54 | 5.08 | 0.041 |
| Interaction (IC-IU)-(CC-CU) |  | MNI |  |  | Peak |  |
| Anatomical location | Voxels | x | y | z | T | $p_{FWE}$ |
| L. Inferior parietal lobule / intraparietal sulcus | 8 | -46 | -46 | -56 | 3.70 | 0.015 |
| L. Anterior intraparietal sulcus | 5 | -36 | -42 | 40 | 3.47 | 0.040 |

CC = cued/congruent, CI = cued/incongruent, UC = uncued/congruent, UI = uncued/incongruent.

Subsequently, we looked at condition-specific activation differences during the execution period depending on the cues presented during the prior delay period. The left aIPS and the left PMd showed significant BOLD signal differences depending on cueing (Fig. 6A and Table 3; cf. Fig. S4 for uncorrected renders). A post-hoc t-test in these areas revealed significantly increased activation in trials where the movement type was left uncued until the ‘Go’ signal, compared with cued trials (*p*FWE<0.05). A whole-brain analysis revealed one further significant voxel in the ventromedial prefrontal cortex. Exploratory correlation analyses revealed that, in the key frontoparietal areas, stronger preparatory activity during cued>uncued trials was associated with lower execution related activity (i.e., cued<uncued) and with faster movement onsets across participants (Fig. S6, but not all of these correlations were significant). Furthermore, an analysis of the loadings on the temporal derivative suggested significantly earlier hemodynamic responses in bilateral frontoparietal areas, but not M1, for uncued > cued movement execution (Fig. S2). Movement execution also depended on cued visuomotor mapping (congruent vs incongruent virtual hand movement) in the left PMd, the left aIPS, and the left anterior insula (Fig. 6B and Table 3; cf. Fig. S4 for uncorrected renders). A post-hoc t-test in these areas revealed significantly increased activation during movements following cued incongruent > congruent visuomotor mapping (*p*FWE<0.05). The main effects overlapped in the left aIPS (conjunction of main effects: x = -38, y = -50, z = 50, peak *F* = 14.57, Fig. S4), but this effect did not reach corrected significance; there were no significant interaction effects. Most of the key brain regions identified in the planning period contrasts (Figs. 3-5) showed substantially stronger BOLD responses during movement compared with planning (Fig. 6C)—with the notable exception of preparation for cued movements under expected incongruence in the left aIPS (CI, cf. Fig. 5), and preparation for incongruent mappings in the right cerebellum (CI and UI, cf. Fig. 4).

**Table 3.** Brain areas showing significant activation differences during movement execution related to prior (cued) movement planning and visuomotor mapping predictions.

| Execution after Movement type cued vs uncued |  | MNI |  |  | Peak |  |
| --- | --- | --- | --- | --- | --- | --- |
| Anatomical location | Voxels | x | y | z | F | $p_{FWE}$ |
| L. Anterior intraparietal sulcus | 12 | -42 | -44 | 50 | 18.88 | 0.020 |
| L. Superior frontal gyrus (PMd) | 7 | -20 | 4 | 70 | 18.69 | 0.021 |
| Ventromedial prefrontal cortex | 3 | 0 | 28 | -6 | 27.74 | 0.047* |
| Execution after Mapping cued congruent vs incongruent |  | MNI |  |  | Peak |  |
| Anatomical location | Voxels | x | y | z | F | $p_{FWE}$ |
| L. Precentral / middle frontal gyrus (PMd) | 14 | -26 | -6 | 50 | 21.59 | 0.017 |
| L. Anterior intraparietal sulcus | 7 | -36 | -50 | 42 | 20.85 | 0.022 |
| L. Anterior insular cortex | 5 | -30 | 24 | 0 | 20.51 | 0.025 |

\* whole-brain analysis.

We also tested for potential ‘expectation violation’ effects observable during movement execution in trials where the virtual hand behaved contrary to the cued expectations (i.e., the cued visuomotor mapping was invalidated). We tested for such responses in brain regions showing increased activity during the preparation for incongruent > congruent mappings, as we assumed this contrast identified brain areas generating visuomotor predictions. Indeed, we found significant (*p*FWE<0.05) surprise effects in the (pre)SMA, the left PMd and PMv, the bilateral IPL/IPS, the left anterior insula, and the right anterior cerebellum (Fig. 6C). An additional whole-brain analysis revealed further surprise effects in the dorsomedial prefrontal cortex, the bilateral inferior frontal gyri, the bilateral angular gyri, the right anterior insula, the right middle temporal and fusiform gyri, the putamen and thalamus, and in the left primary sensory and motor cortex (Fig. S5). There were no significant differences in surprise-related activations between conditions (i.e., no voxels showed a significant main or interaction effect). The converse contrast (expected > unexpected visuomotor mapping) yielded no significant activations either.

Finally, following the observation that the movement cueing effects in our left-hand experiment were strongly left-lateralized in the IPS, six of our original study’s participants were invited to a separate scanning session, in which they repeated a right-hand version of the task. These results are reported in Figure S7. Most importantly, as in the main experiment, the preparation of cued > uncued movements was associated with stronger BOLD signal in bilateral frontoparietal areas and the V5. Within the left, compared with the right IPS, the effects were more extensive (spanning to anterior portions of the IPS only in the left hemisphere) and stronger (all peak T-values were higher in the left > right PPC). Again, the preparation of movements with an expected incongruent > congruent visuomotor mapping was associated with stronger BOLD signal in the cerebellum and in bilateral frontoparietal areas. Contrasting the execution of uncued > cued movements likewise revealed activation differences corresponding to those in the main experiment; and again stronger left-hemispheric effects in the PPC. In sum, the pattern of results obtained during *right-hand* planning and movement replicated our main results; together, our results suggest that motor planning, in our design, predominantly engaged the left IPS for either hand side.

## Discussion

In our non-spatial delayed movement task with altered visuomotor mapping, delay period activity in the premotor cortex (PMd, PMv, and the SMA) and the PPC (IPS and SPL) was increased if one of two alternative movements was cued ahead of execution. This supports the general association of those regions with motor planning (see Introduction) and aligns with frontoparietal BOLD signal increases observed during cued reach planning (Fernandez-Ruiz et al., 2007; Lindner et al., 2010; Gertz & Fiehler, 2015; Pilacinski et al., 2018). The PMd and aIPS showed the inverse pattern during execution (relatively higher BOLD when executing uncued movements), as similarly observed in previous uncued vs cued effector selection tasks (Bernier et al., 2012). We propose this indicates the fast establishment of an ad-hoc motor plan, which our participants did not already prepare during the delay (cf. Ames et al., 2014; Pilacinski et al., 2018). In other words, our participants seemed to simply wait for the execution prompt in the uncued trials, instead of preparing both possible movements during the delay. Whether (or perhaps rather: when) the motor system plans actions serially, one at a time, or multiple potential plans simultaneously, is a key question of the current motor control literature—with evidence for both cases (e.g., Dekleva et al., 2018; Cisek & Kalaska, 2002, 2005; Cui & Andersen, 2011). The results of such studies depend, among other things, on the kind of decision, the number of alternatives, and the kind of movement used in each case. For instance, “nonspatial motor decisions” – like in our task – may be processed more serially than others (Cui & Andersen, 2011); intransitive actions may also be processed differently than goal-directed actions (see below). In our case, the frontoparietal BOLD pattern and the behavioral reaction time differences clearly indicated the establishment of one specific movement plan during the delay of cued trials, rather than parallel planning of both possible movements (and subsequent selection of the prompted movement) during the delay of uncued trials.

Our second key finding was the relatively higher cerebellar activity during the preparation for incongruent > congruent visuomotor mappings. In addition to its higher amplitude, the (right) cerebellar hemodynamic response was relatively delayed (i.e., it peaked later) and wider in those cases. This suggests that, when the associated visual movement feedback was expected to be incongruent (i.e., inverted) while otherwise identical muscle (hand) movements were planned, cerebellar neural activity was elevated and more sustained. Thereby, in the right cerebellum, the amplitude of the BOLD response during preparation for incongruence even exceeded that observed during actual movement execution. Note that the incongruent virtual hand movements were predictable in principle from (planned) motor signals, just like the congruent ones, because they reflected the (inverted) sensor values from the data glove; i.e., their kinematics were also controlled by the participant’s hand movements. Taken together, we propose that the associated stronger cerebellar delay activity likely indicated an anticipatory update of forward predictions of visual movement feedback—of “What my hand movement will look like”—specifically in those conditions where the expected visual hand movement feedback violated the participants’ life-long learned visuomotor associations (i.e., of “congruence” between executed hand movements and their visual feedback). Conversely, the ‘natural’ predictions of visuomotor congruence could have been more easily and more quickly generated, requiring less computational resources (cf. Kavroulakis et al., 2022). That the preparation for incongruent mappings increased cerebellar BOLD regardless of whether a specific movement was planned or not (as indicated by the significant conjunction contrast of preparation for incongruence > congruence across cued and uncued movements) suggests a general update of the effector’s forward model to a nonstandard visuomotor mapping; i.e., not only of those predictions associated with the specific movement selected for execution.

It has long been proposed that the cerebellum implements forward models (Miall et al., 1993; Wolpert et al., 1998). Lobule VI, specifically, responds to unpredicted sensory movement feedback (Blakemore et al., 1998, 2001; Kilteni et al., 2023), including visuomotor incongruencies (Leube et al., 2003; Limanowski et al., 2017; Van Kemenade et al., 2019). During sensorimotor adaptation, cerebellar activity may indicate trial-to-trial updates of forward models by sensory prediction errors (Shadmehr & Krakauer 2008; Izawa et al., 2012; see Tseng et al. 2007; Synofzik et al., 2008; Donchin et al. 2012; Tzvi et al. 2022 for visuomotor adaptation, specifically). Our results suggest that cerebellar predictions —likely informing state estimates about visuomotor mapping—can be updated ahead of, i.e., without movement execution, and thus without any associated sensory prediction errors. This supports the hypothesized generation of forward sensory predictions during action planning (see Introduction). The other main regions showing increased activity during the preparation for incongruence were the PMd, which has been shown to implement adjustments to “nonstandard sensorimotor mapping” (Wise et al., 1996); and the SMA. The SMA (the dorsomedial frontal cortex in general) is involved in processing conflicts during action selection (Taylor et al., 2007; Duque et al., 2013), error learning (Ulllsperger et al., 2014), and visuomotor adaptation (Grafton et al., 2008; Nahab et al., 2011; Limanowski et al., 2017; Ohata et al., 2020). Furthermore, electrophysiological recordings suggested that the “readiness potential”, which likely originates from the SMA, may also represent a prediction of (voluntary) action feedback prior to execution—much like proposed by the efference copy concept (Reznik et al., 2018; Pinhero et al., 2020; Ody et al., 2023). Both premotor areas could therefore have contributed to forward modeling (although the clearest predictive effects were observed in the cerebellum), or more generally anticipated the upcoming conflict (here: a nonstandard visuomotor mapping).

Thirdly, and crucially, the left aIPS showed a significant interaction effect, characterized by relatively increased BOLD during the planning of a pre-specified (cued > uncued) movement under an expected nonstandard (incongruent > congruent) visuomotor mapping. While several fronto-parietal areas showed overlapping activations by both main effects (in line with suggestions that these regions may be involved in more than one type of computation for motor control, cf. Scott, 2012), only the left aIPS/IPL showed a significant interaction effect. Thus, movement planning in the aIPS was more metabolically demanding when nonstandard>standard visuomotor mappings were expected. We propose that, while cerebellar activity could be linked to forward model updates, the interaction effect in the aIPS indicated a different computation; namely, the *integration* of the respective updated (cerebellar) forward predictions of nonstandard visual body movement feedback with the state estimate used for movement planning. The fact the interaction was confined to the anterior part of the IPS fits with this region’s proposed role of a specifically “body-related” state estimator (Medendorp & Heed, 2019; Kuang et al., 2015). It also aligns with previous demonstrations of the aIPS’s key role in adaptive visuomotor control and upper limb state estimation (Desmurget et al., 1999; Brass et al., 2001; Ehrsson et al., 2004; Grefkes & Fink, 2005; de Lange et al., 2006; Zimmermann et al., 2012; Limanowski & Blankenburg, 2016, 2017; Limanowski, 2022). Generally, it supports the notion of the PPC as a high-level motor control region (Andersen and Buneo 2002; Blakemore & Sirigu, 2003).

Our results highlighted specifically the *left* aIPS, irrespective of the hand side used (i.e., analogous effects and topography in our main, left-hand experiment and a right-hand replication, see Supplementary Material). This speaks to previous findings: The PPC represents actions from contra- and ipsilateral hands (Gallivan et al., 2013; Turella et al., 2020); and the left (but not right) PPC has been found activated during the selection, preparation, and execution of hand actions irrespective of hand laterality (Schluter et al., 2001; Rushworth et al., 2001; Grafton et al., 2002; Hardwick et al., 2013). The left PPC has also been identified as playing a key role in motor imagery, visuomotor adaptation, and the visuoproprioceptive integration it entails, for the left and right hand (Bonda et al., 1995; Servos et a., 2002; de Lange et al., 2006; Mutha et al., 2011; Hétu et al., 2013; Limanowski & Blankenburg, 2017). Notably, left PPC lesions have been associated with ideamotor apraxia (Haaland et al., 2000); fMRI results correspondingly suggest a “left-lateralized praxis network”, centered on the PPC, which plans actions with either hand (Bohlhalter et al., 2009). Specifically, intransitive actions (i.e., without object interaction) may be more strongly left-lateralized than transitive ones (Króliczak & Frey, 2009). Our results, obtained with such intransitive actions, fit very well into this view; moreover, they show that the left PPC is essential for integrating visual movement predictions with the respective motor plans.

Two regions showing a main effect of planning but not visuomotor incongruence were the PMv and the V5. The PMv is clearly linked to upper limb motor control (Graziano, 1999); the V5 activation could indicate the generation of visual postural or motion templates (Zimmermann et al., 2012, 2016; Yon et al., 2018), perhaps as ‘simulated’ sensory input for state estimation. Finally, the cerebellum, the SMA, and several other regions that increased activity during the preparation for incongruent visuomotor mappings showed surprise related responses. Thus, brain regions updating predictions also responded to the respective prediction errors. That surprise responses did not differ between congruent and incongruent mappings suggests that the forward model’s predictions were, generally, successfully updated in response to the cues. Further surprise responses were located in medial and inferior frontal areas associated with general error detection (Ullsperger et al., 2014; cf. Quirmbach & Limanowski, 2022) and in temporoparietal areas implied in visuomotor mismatch detection (Leube et al., 2003; Balslev et al., 2005; Farrer et al., 2008; van Kemenade et al., 2017, 2019; Limanowski et al., 2017). That not all of the ‘predictive’ regions showed surprise responses, and vice versa, can potentially help refine neuroanatomical models of visuomotor adaptation (Synofzik et al., 2006; Grafton et al., 2008).

Together, our results support the idea of forward sensory prediction during action planning, and associate the underlying computations mainly with the cerebellum and the PPC. This opens up interesting avenues for future work, for instance, using brain stimulation. Thus, previous work has shown that transcranial magnetic or direct current stimulation of the aIPS or the cerebellum directly influenced adaptive motor control (Desmurget et al., 1999; Yavari et al., 2016; Cao et al., 2017; cf. Weightman et al., 2023). Stimulation of frontoparietal and cerebellar areas improved sensorimotor delay detection and recalibration in schizophrenic patients, which presumably suffer from deficits in forward modeling (Straube et al., 2020; Schmitter & Straube, 2024; cf. Voss et al., 2010). One question following from our work is whether such cerebellar or posterior parietal manipulations would similarly affect the updating and integration of sensory predictions during action *planning*. Future studies could also use multivariate approaches to reveal the encoding of more abstract action planning related information such as, e.g., different sensory content representations (cf. Ariani et al., 2015, 2018; Gallivan et al., 2013; Yon et al., 2018; Turella et al., 2020). Our study raises further questions: One limitation of our design were the relatively frequent catch trials that more likely followed invalid (surprising) than valid movements. Participants could thus have associated surprising movements with subsequent button presses, introducing a bias by movement preparation, initiation, or reward expectations. Thus, some surprise effects, e.g. in motor areas, could partly reflect anticipatory or more extensive movements of the responding (right) hand; which needs to be clarified by future work. However, this did not affect the between-condition contrasts of the movement (or preparation) periods, as surprise and catch trials were balanced across conditions. Furthermore, we cannot exclude potential eye movement influences on our results (cf. Gorbet & Sergio, 2016). However, our non- spatial movement task and the instructed central fixation rendered systematic eye movements useless; and our control experiment suggested only minimal differences in gaze behavior between conditions.

To conclude, our results suggest visual feedback prediction is an inherent component of motor planning; and a key role of the cerebellum and the left aIPS in the underlying forward predictions and integrative processes, respectively.

## Acknowledgments

This work was funded by the German Research Foundation (DFG, Deutsche Forschungsgemeinschaft) as part of Germany’s Excellence Strategy – EXC 2050/1 – Project ID 390696704 – Cluster of Excellence “Centre for Tactile Internet with Human-in-the-Loop” (CeTI) of Technische Universität Dresden. JL was supported by a Freigeist Fellowship of the VolkswagenStiftung (AZ 97-932). We thank Gesche Vigh for help with implementing the virtual reality.

