## Supplementary Material for "Visuomotor prediction during action planning in the human frontoparietal cortex and cerebellum"

**Authors:** Felix Quirmbach[1,2] and Jakub Limanowski[2,3]
[1]Faculty of Psychology, Technical University of Dresden, 01069 Dresden, Germany
[2]Center for Tactile Internet with Human-in-the-Loop, Technical University of Dresden, 01187 Dresden, Germany
[3]Institute of Psychology, University of Greifswald, 17486 Greifswald, Germany

---

#### Fig. S1: Eye tracking control experiment

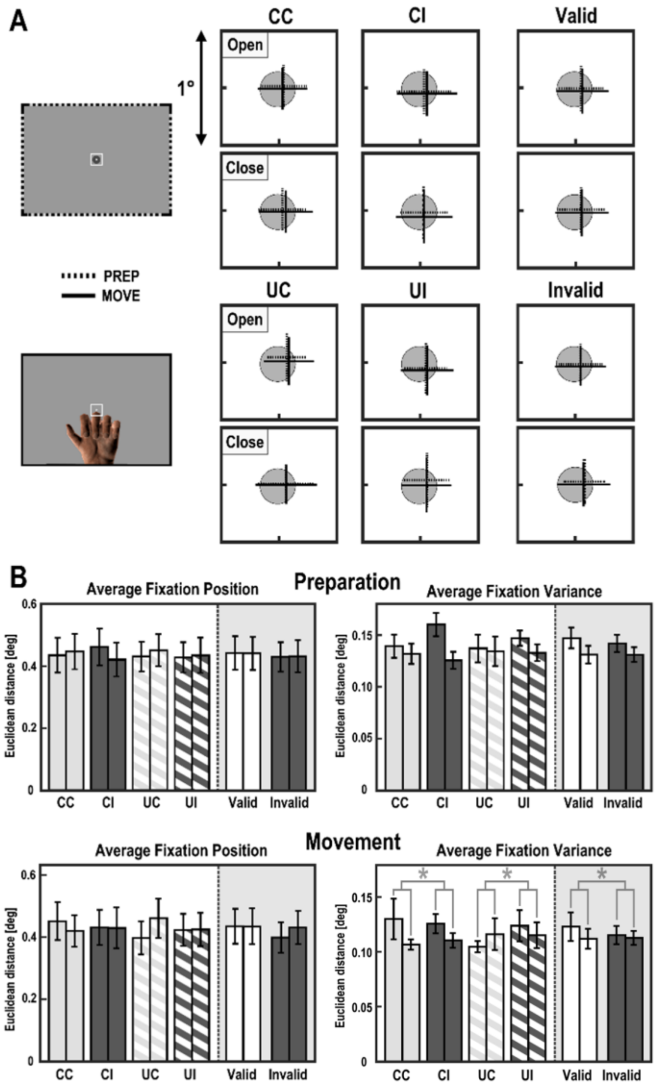


**A:** A separate eye tracking experiment (*N*=6 of the original study participants, without scanning) showed that average fixation position did not significantly differ between conditions during the preparation or execution periods. The right plots show, for open and close movements separately, a small section of the screen (marked by the white squares in the screenshots) centered on the fixation dot (grey). The crosses mark the respective average fixation, with horizontal and vertical standard deviation between subjects during the preparation period (dotted bars) and the execution period (solid bars CC = cued/congruent, CI = cued/incongruent, UC = uncued/congruent, UI = uncued/incongruent. **B:** Average fixation position and average fixation variance in the individual conditions for both the preparation and movement phase, in Euclidean distance from the fixation point, with associated standard errors of the mean. Conditions (CC, CI, UC, UI) like above, pooled across valid/invalid conditions (white background) or for validly/invalidly cued trials, pooled across conditions (dark background). For each pair of bars, the first bar corresponds to the "open" hand movement, the second bar to the "close" movement. Average fixation position did not significantly differ depending on any experimental factor (a significant three-way-interaction (visuomotor mapping x cueing x movement type) was significant for both preparation and movement phases, but difficult to interpret). There was a significant, but relatively weak main effect of visuomotor mapping on fixation variance during the execution period ($F_{(1,5)} = 7.25$, $p = 0.043$, denoted by grey asterisks in the bottom right bar plot); however, of all post-hoc t-tests across individual conditions, only one reached significance (UI valid, open > UC valid, open). Importantly, fixation variability was in itself very small: the average standard deviation of any condition was <0.17 degrees visual angle, whereas the hand movements spanned 10.6 degrees. Thus, the eye tracking results suggested that the results obtained from our fMRI contrasts were likely not biased by systematic differences in gaze behavior.

### Fig. S2: Analysis of loadings on the temporal and dispersion derivatives

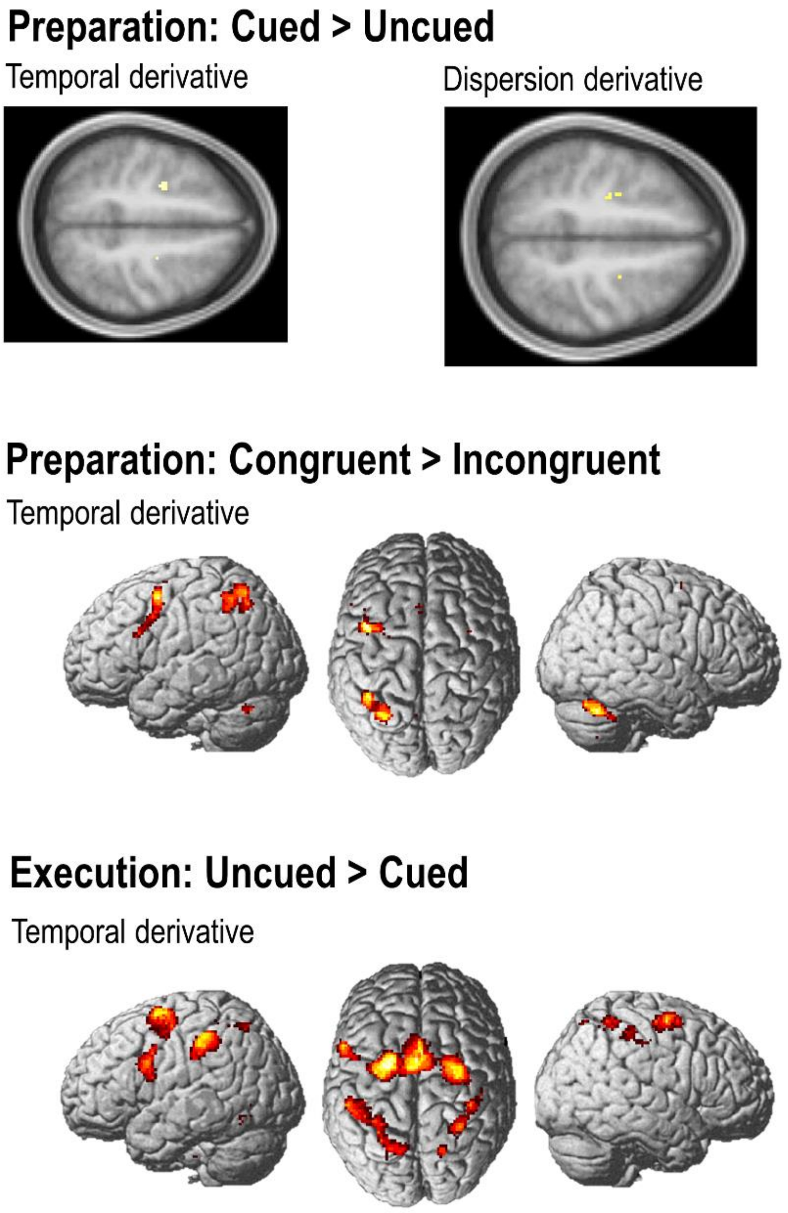


While the main aim of our fMRI analysis was to look at relative BOLD signal amplitude, indicating varying metabolic demand, predictive processes may also be inferred from differences in the relative latency and width of the BOLD response (Kavroulakis et al., 2022). Therefore, we looked for differences in the conditional loadings on the temporal and dispersion derivatives of the conditional regressors included in our model (cf. Friston et al., 1998). **A:** For BOLD responses related to the *preparation* cues, an analysis of the loadings on the temporal derivative revealed significantly earlier responses related to the preparation of cued > uncued movements in the bilateral PMd ($p_{FWE}$<0.05; no significant effects in the converse contrast). However, these regions fell outside of the areas showing an amplitude main effect. Furthermore, we found significantly earlier responses related to the preparation of congruent > incongruent mappings in the bilateral cerebellum (see Fig. 4), with the strongest effect again in the right cerebellum; and furthermore in the left IPS and PMv, the bilateral PMd, and the SMA ($p_{FWE}$<0.05; no significant effects in the converse contrast). There was no significant interaction effect on the temporal derivative; one voxel in the cerebellum showed a significant interaction effect in terms of dispersion, but this voxel fell outside of any area activated by the amplitude analysis (x = -24, y = -42, z = 52). **C:** For the *movement* period, an analysis of the loadings on the temporal derivative suggested significantly earlier onsets of BOLD responses in several bilateral frontoparietal areas for uncued > cued movements (no significant converse effects). Notably, there were no timing differences in the contralateral M1; suggesting these temporal shifts were not mainly related to movement production itself. Here, it has to be remembered that the onsets of the regressors were not centered on the actual movement onsets but on the appearance of the virtual hand (in order to primarily capture visuomotor comparisons, see Methods). In this light, the above differences should only carefully be interpreted. Yet, they could indicate the generation of an ‘ad hoc’ motor plan prior to the actual movement execution (i.e., after the directional cue but before the ‘GO signal’) in the uncued conditions (whereas in the cued conditions no or less of such activity would be needed, as the motor plan had already been established). See Discussion. There were significantly earlier BOLD onsets in the cerebellum for the execution after cued congruent > incongruent mappings (no converse effects), but these only spanned 1 voxel and fell outside of the cerebellar areas engaged by predictive processes during the preparation period.

### Fig. S3: Conjunction and interaction of main effects during the delay period

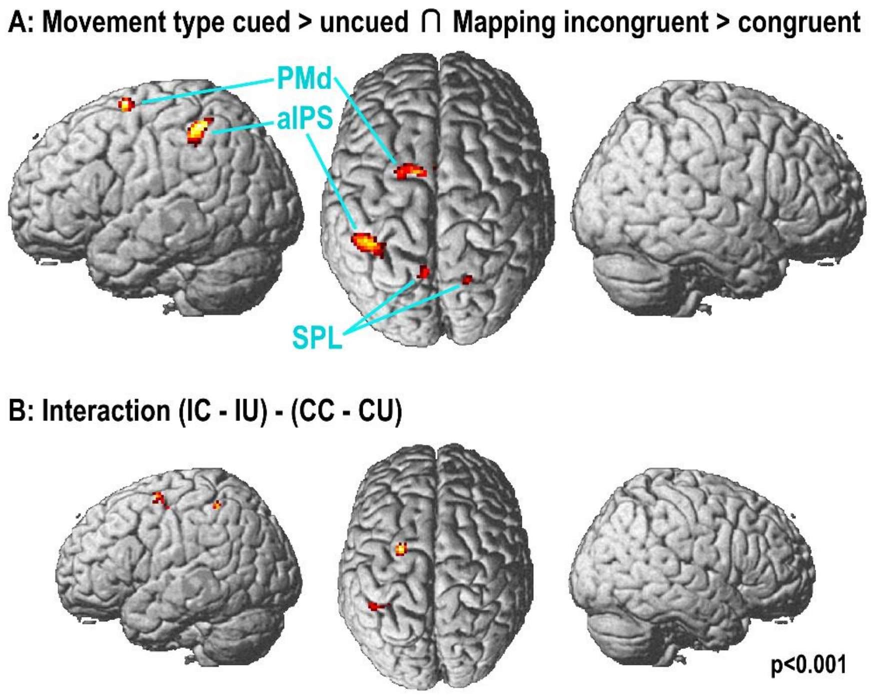


**A:** Render showing voxels with significantly ($p_{FWE}$<0.05) stronger BOLD during the preparation of cued > uncued movements and during the preparation for incongruent > congruent visuomotor mappings; i.e., a null conjunction of the main effects shown in Figs. 3A and 4: (CC+CI)-(UC+UI) ∩ (CI+UI)-(CC+UC). **B:** Uncorrected ($p$<0.001) activations obtained from the interaction effect in Fig. 5; showing that the effect was localized to the left aIPS and an additional peak in the left PMd (which, however, did not reach corrected significance; x = -26, y = -4, z = 58, T = 3.71).

### Fig. S4: Uncorrected activation differences during the movement period.

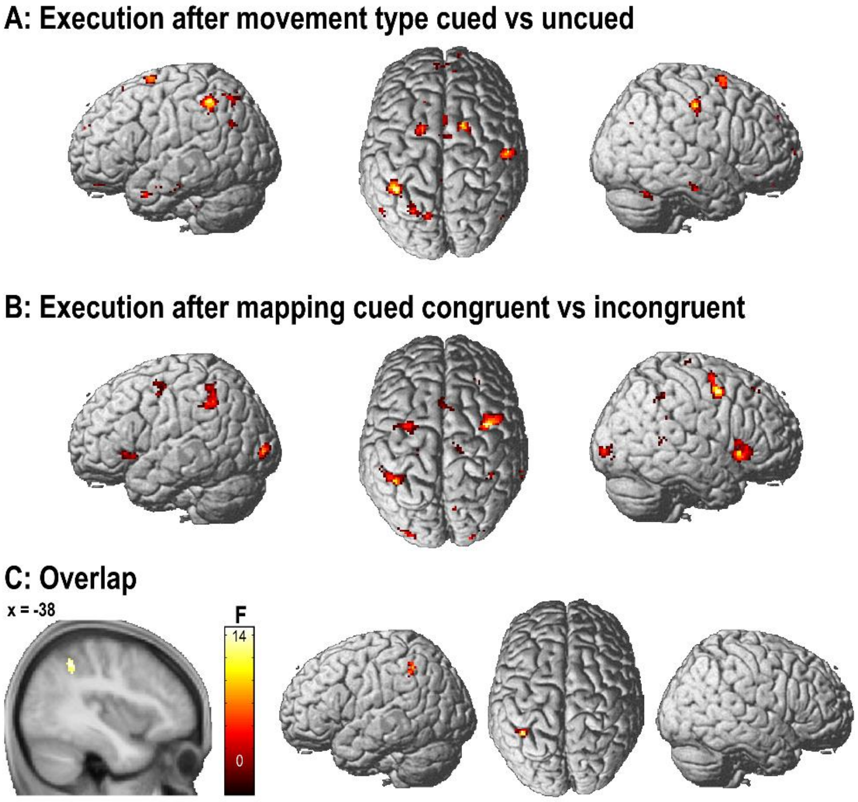


**A:** Renders of voxels showing BOLD signal differences (F-test at *p*<0.001, uncorrected) during movement execution in trials where the movement type had been cued in advance versus when it was left ambiguous (uncued); related to Fig. 5A. **B:** Renders of voxels showing BOLD signal differences (F-test at *p*<0.001, uncorrected) during movement execution in trials where congruent vs incongruent visuomotor mapping had been cued; related to Fig. 5B. **C:** Voxels in the left aIPS showing an overlap of both main effects during the movement period (conjunction contrast, *p*<0.001, uncorrected).

### Fig. S5 Whole-brain surprise effects

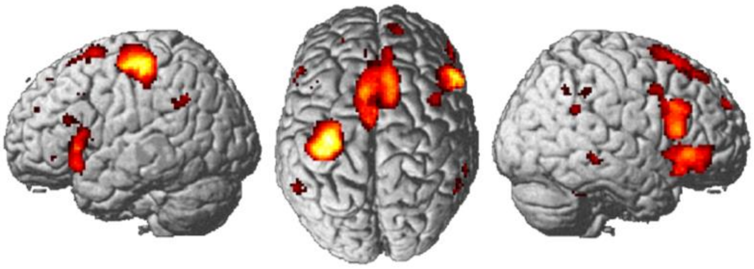

Render of significant ($p_{FWE}$<0.05) activations during unexpected > expected visuomotor mappings in a whole-brain analysis. Related to Fig. 6C.

### Fig. S6: Exploratory correlation analysis

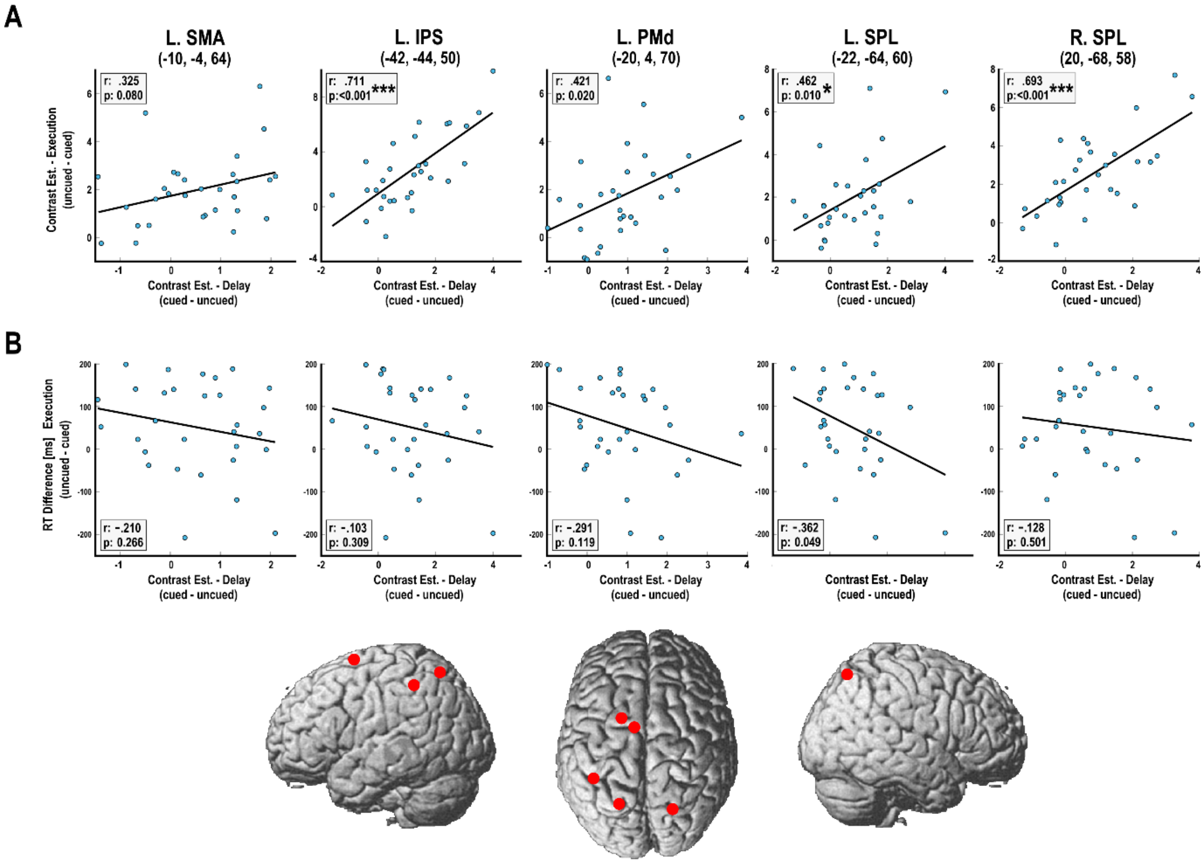


In our main analysis, we observed increased activation in similar (frontoparietal) areas during the preparation phase for trials with movement type cued > uncued (cf. Fig 3, Table 1) and during the execution phase for trials with uncued > cued movement type (cf. Fig 6, Table 3). We also observed earlier hand movement onset (reaction time, RT) when movement type was cued vs left ambiguous (cf. Fig. 2A, S1). Following a suggestion from one of our reviewers, we tested – within the key frontoparietal areas identified in our main contrasts – for correlations between activation levels between the experimental phases, and a potential correlation between activation in the preparation phase and behavior i.e. movement onset (reaction time). For each participant and region of interest, we extracted the first-level contrast estimate from the group peak voxel for the contrast cued > uncued movement type in the *preparation* phase, and from the contrast *execution* after movement type uncued > cued. As our key regions of interest, we selected voxels showing significant BOLD signal differences during execution depending on cueing (i.e., left IPS and PMd; cf. Fig. 6, Table 3), as well as prominent regions of activation during the preparation phase for contrast cued > uncued (i.e., left SMA and bilateral SPL; cf. Table 1). As our behavioral measure, we calculated the difference of average movement reaction times between trials with uncued vs cued movement type, i.e., how much longer it took participants to initiate a hand movement when the movement type was left uncued. Correlation was determined by calculating the Pearson correlation coefficient between measures, with the significance level adjusted for multiple comparisons (in the Figure, significant correlations are marked via asterisk: *$p$<0.05/5=0.01, ***$p$<0.001 = <0.0002). **A:** Plots showing the relationship between preparation and execution phase activity. Increased activation for cued > uncued trials during the preparation phase was generally associated with increased activation for uncued > cued trials during the execution phase in all five regions. Rho-values ranged between .33 and .71, with significant positive correlation (p<0.01) in the left IPS and bilateral SPL and a trend towards significance in left SMA and PMd. In other words, the stronger the activation increase during preparation for cued movements, the lower the activation to those trials (compared with uncued movements) during execution. Tentatively, this correlation supports our interpretation of average activity differences; i.e., that pre-cueing specific movements led to activity increase per establishing a motor plan, whereas for uncued movements, no such preparation was evident – and instead an ad-hoc motor plan was calculated upon execution. **B:** Plots showing brain-behavior correlations. We found hints towards an association between increased preparatory activity and hand movement onsets; i.e., relatively stronger preparatory activity during cued > uncued trials was associated with relatively faster movement onsets (cued < uncued). Tentatively, this supports our idea that pre-cueing a specific movement established a motor plan, which was initially lacking when uncued movements were prompted. However, while rho-values were generally negative, these correlations did not reach significance after correction in any region.

### Fig. S7: Right-hand fMRI replication experiment.

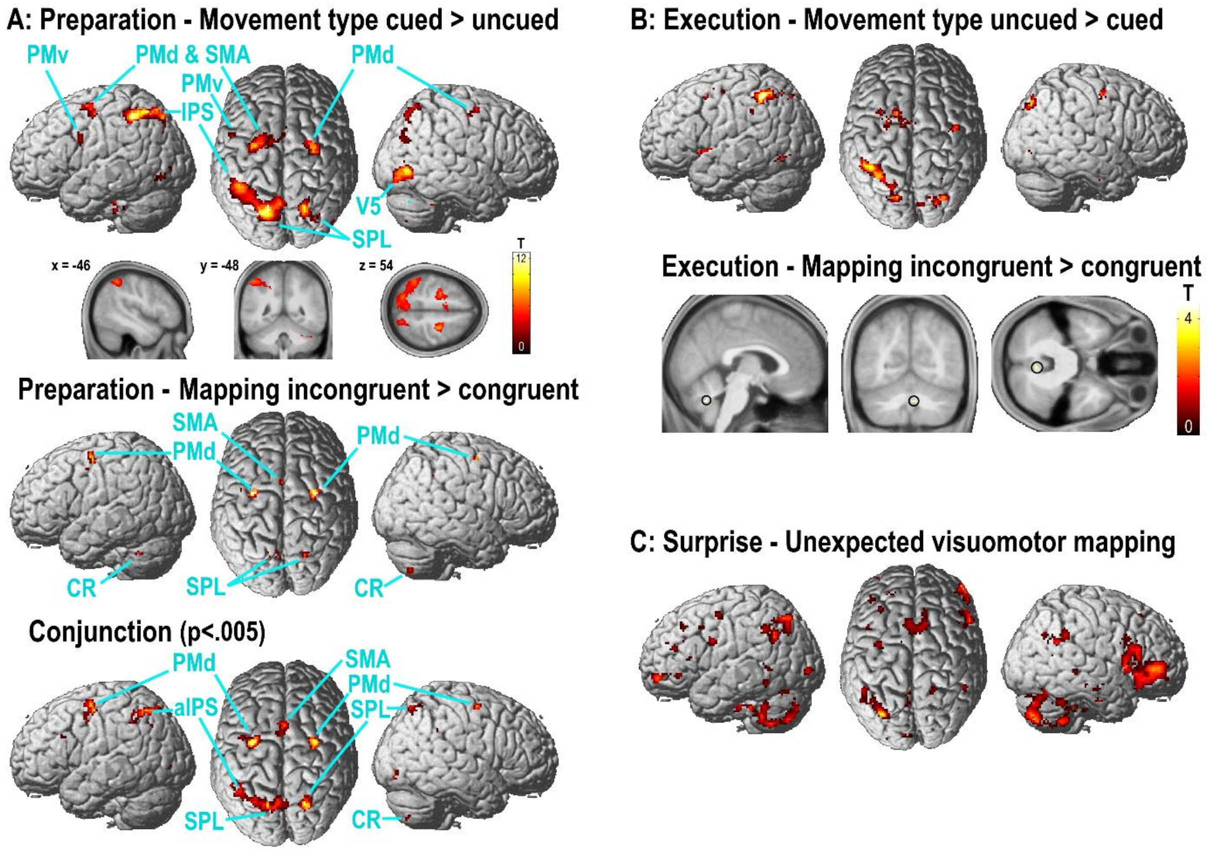


*N*=6 of the original study participants repeated the main experiment with right hand movements. The renders show group-level activations ($p$<0.001, uncorrected) obtained from the respective contrasts. The conjunction contrast was significant at $p$<0.001, but is displayed at $p$<0.005 to better show the spatial extent of left > right-hemispheric PPC effects. **A:** Preparation period (cf. Figs. 3A,B); The interaction effect did not reach significance but was present, among other regions, in the left but not right IPL (x = -48, y = -58, z = 50, T = 3.13, p<0.005). **B:** execution period (cf. Fig. 5); **C:** surprise effect (cf. Fig. 5C).
